# Freeze-frame imaging of synaptic activity using SynTagMA

**DOI:** 10.1101/538041

**Authors:** Alberto Perez-Alvarez, Brenna C. Fearey, Ryan J. O’Toole, Wei Yang, Ignacio Arganda-Carreras, Paul J. Lamothe-Molina, Benjamien Moeyaert, Manuel A. Mohr, Lauren C. Panzera, Christian Schulze, Eric R. Schreiter, J. Simon Wiegert, Christine E. Gee, Michael B. Hoppa, Thomas G. Oertner

**Affiliations:** Institute for Synaptic Physiology, University Medical Center Hamburg-Eppendorf, Hamburg, D-20251 Germany; Department of Biological Sciences, Dartmouth College, Hanover, NH, 03755 USA; HHMI, Janelia Farm Research Campus, Ashburn VA, 20147 USA; Ikerbasque, Basque Foundation for Science, Bilbao, Spain; Dept. of Computer Science and Artificial Intelligence, Basque Country University, San Sebastian, Spain; Donostia International Physics Center (DIPC), San Sebastian, Spain; Research Group Synaptic Wiring and Information Processing, University Medical Center Hamburg-Eppendorf, Hamburg, D-20251 Germany

## Abstract

Information within the brain travels from neuron to neuron across billions of synapses. At any given moment, only a small subset of neurons and synapses are active, but finding the active synapses in brain tissue has been a technical challenge. To tag active synapses in a user-defined time window, we developed SynTagMA. Upon 395-405 nm illumination, this genetically encoded marker of activity converts from green to red fluorescence if, and only if, it is bound to calcium. Targeted to presynaptic terminals, preSynTagMA allows discrimination between active and silent axons. Targeted to excitatory postsynapses, postSynTagMA creates a snapshot of synapses active just before photoconversion. To analyze large datasets, we developed software to identify and track the fluorescence of thousands of individual synapses in tissue. Together, these tools provide an efficient method for repeatedly mapping active neurons and synapses in cell culture, slice preparations, and *in vivo* during behavior.

## MAIN

The physical changes underlying learning and memory likely involve alterations in the strength and/or number of synaptic connections. On the network level, neuroscience faces an extreme ‘needle in the haystack’ problem: it is thought to be impossible, in practice, to create a map of all synapses that are active during a specific sensory input or behavior. Although excellent genetically-encoded sensors for calcium, glutamate and voltage have been developed^1–3^, and two-photon laser-scanning microscopy enables monitoring the activity of neurons and even single synapses in highly light-scattering brain tissue^4^, the tradeoff between spatial and temporal resolution makes it impossible to simultaneously measure fluorescence in the thousands of synapses of even a single pyramidal neuron. Most functional imaging experiments are therefore limited to cell bodies, i.e. low spatial resolution^5^, or monitor the activity of a few synapses within a single focal plane^6^. Multi-beam scanning designs have been proposed, but due to scattering of emitted photons, they do not produce sharp images at depth^7,8^. Projection microscopy can be a very efficient approach^9^, but only in situations where the fluorescent label is restricted to one or very few neurons. In general, the need to choose between high temporal or high spatial resolution limits the information we can extract from the brain with optical methods.

A strategy to overcome this limit is to rapidly ‘freeze’ activity in a defined time window and read it out at high resolution later. The Ca^2+^-modulated photoactivatable ratiometric integrator CaMPARI undergoes an irreversible chromophore change from green to red when the Ca^2+^ bound form is irradiated with violet (390-405 nm) light^10,11^. CaMPARI has been successfully applied to map the activity of thousands of neurons in zebrafish, *Drosophila*, and in mouse^10,12–14^. As CaMPARI was designed to diffuse freely within the cytoplasm, it does not preserve subcellular details of Ca^2+^ signaling. By anchoring CaMPARI to either pre- or postsynaptic compartments, we developed tools that mark active synapses in short time windows defined by violet illumination. Three steps were necessary to create a Synaptic Tag for Mapping Activity (SynTagMA): 1) We improved the brightness and conversion efficiency of CaMPARI^11^; 2) We targeted CaMPARI2 (F391W_L398V) to either presynaptic boutons by fusing it to synaptophysin (preSynTagMA) or to the postsynaptic density by fusing it to an intrabody against PSD95^15^ (postSynTagMA); 3) We developed an analysis workflow that corrects for chromatic aberration, tissue displacement (warping) and automatically finds regions of interest (i.e. postsynapses or boutons) to quantify green and red fluorescence. Bouton-localized preSynTagMA allowed us to distinguish active and inactive axons. Spine-localized postSynTagMA allowed us to map the extent of action potential back-propagation into the large apical dendritic tree of hippocampal pyramidal cells. Following repeated sparse activation of Schaffer collateral axons, postSynTagMA marks a small subset of synapses on CA1 pyramidal cells as highly active. PostSynTagMA is not only useful to analyze the organization of inputs on spiny pyramidal cells, but works equally well for interneurons where most synapses are formed on the dendritic shaft. A fraction of postSynTagMA is sequestered to the nucleoplasm, generating a useful label to identify neurons that are active during a specific behavior. As an application example, we photoconverted active CA1 pyramidal cells during reward collection in a spatial memory task. In summary, the key advantage of SynTagMA compared to acute calcium sensors is the greatly extended read-out period, allowing three-dimensional scanning of relatively large tissue volumes at cellular or at synaptic resolution.

## RESULTS

### Creating and characterizing a presynaptic marker of activity

To visualize activated presynaptic boutons, we fused the Ca^2+^-modulated photoactivatable ratiometric integrator CaMPARI^10^ to the vesicular protein synaptophysin (sypCaMPARI). In cultured hippocampal neurons, sypCaMPARI showed a punctate expression pattern along the axon, indicating successful targeting to vesicle clusters in presynaptic boutons (**Fig. 1a**). SypCaMPARI fluorescence decreased after stimulating neurons to evoke trains of action potentials (APs), saturating at 50 APs and slowly returning to baseline (**Fig. 1b-d**). This Ca^2+^-dependent dimming, which is a known property of CaMPARI ^10^, provides a low-pass-filtered read-out of ongoing neuronal activity. Indeed, the signal-to-noise ratio was sufficient for detecting single APs in single boutons when 30 sweeps were averaged, suggesting a high sensitivity of localized sypCaMPARI for Ca^2+^ influx (**Fig. 1c**). The Ca^2+^-bound form of CaMPARI is irreversibly photoconverted from a green to a red fluorescent state by 405 nm irradiation^10^. Repeatedly pairing electrical field stimulation with 405 nm light flashes increased the red-to-green fluorescence ratio (R/G) of sypCaMPARI boutons in a stepwise fashion (**Fig. 1e and f**). Blocking action potential generation with tetrodotoxin (TTX) strongly reduced photoconversion, indicating that spike-induced Ca^2+^ influx through high-threshold voltage-activated Ca^2+^ channels was necessary for efficient conversion. Similar to the dimming response, the amount of sypCaMPARI photoconversion depended on the number of APs (**Fig. 1g**), suggesting that the R/G ratio can be interpreted as a lasting and quantitative record of axonal activity rather than just a binary signal.

**Figure 1.**
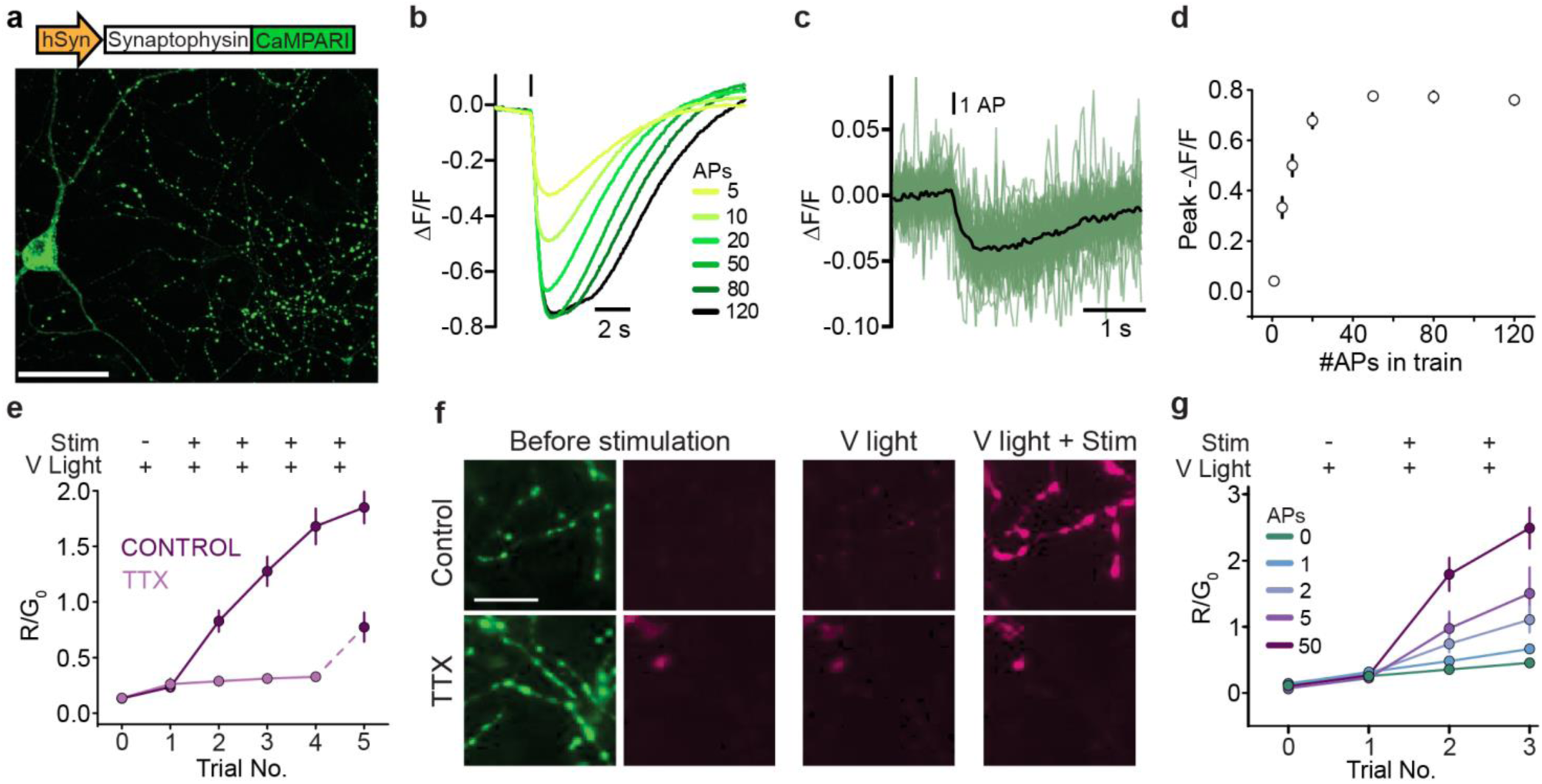
Synaptophysin-fused CaMPARI marks active presynaptic terminals. **(a)** Representative image of cultured rat hippocampal neurons expressing sypCaMPARI. Note the clear punctate labeling of axonal boutons. Scale bar 50 µm. **(b)** Average fluorescence response of sypCaMPARI boutons (green channel emission) to varying numbers of action potentials (APs) evoked at 50 Hz (n = 6 neurons). **(c)** Trial-averaged responses to 30 single APs (green, n = 57 boutons). Black line is the average representative of n = 3 neurons. **(d)** Plot of the maximum ΔF/F versus number of APs from the experiments in **b** and **c** (n = 6 cells, 317 synapses). **(e)** Plot of initial red to green ratio of boutons expressing sypCaMPARI (R_0_/G_0_, V light -, Stim -), after photoconverting violet light alone (405 nm, 20 flashes x 1 s, 0.1 Hz, 10.8 mW cm^-2^, V light +, Stim -) and after simultaneous stimulation with trains of 50 APs at 50 Hz (V light +, Stim +, repeated 4 times). The experiment was performed in the absence or presence of 3 µM tetrodotoxin to block action potentials (control: n = 8 cells; TTX: n = 7 cells). Note that after washing out TTX, the R/G_0_ ratio (V light + Stim +, trial 5) increased to the same extent as the first instance in control neurons (trial 2). **(f)** Representative red (magenta, Trial 0, Trial 1, Trial 3) and green (green, Trial 0) images of boutons from the experiment in **e**. The scale bar 20 µm applies to all panels in **f. (g)** The amount of photoconversion (R/G_0_) in a similar experiment as **e** but varying the number of APs in a 50 Hz train (V light 405 nm, 20 flashes x 1 s, 0.1 Hz, 54.1 mW cm^-2^, Stim 20 bursts at 50 Hz). Data are mean ± SE (0 AP: n = 4 cells, 1 AP: n = 4, 2 APs: n = 5, 5 APs n = 5, 50 APs n = 7). 50 AP fold increase was statistically different from all other stimulation conditions using a one-way ANOVA with Tukey’s post-hoc comparison (p < 0.05).

Ca^2+^-dependent dimming of sypCaMPARI peaked about 1 s after stimulation and returned to baseline after 3-15 s (**Fig. 1b, c**). To determine the photoconversion time window, multiple delays were tested in single experiments using a digital mirror device (**Fig. 2a**). Photoconversion of sypCaMPARI was not efficient when light pulses (100 ms duration) were coincident with stimulation onset (**Fig. 2b, c**). Rather, maximal photoconversion occurred when violet light was applied 2-5 s after stimulation. The photoconversion window of sypCaMPARI extended to at least 10 s after stimulation, outlasting the dimming response to 5 APs (**Fig. 1b**). Some photoconversion also occurred in the absence of stimulation or in the presence of TTX (**Fig. 1e, g** and **Fig. 2b, c**). The long temporal window and activity-independent photoconversion are both undesirable traits that limit the utility of sypCaMPARI.

**Figure 2.**
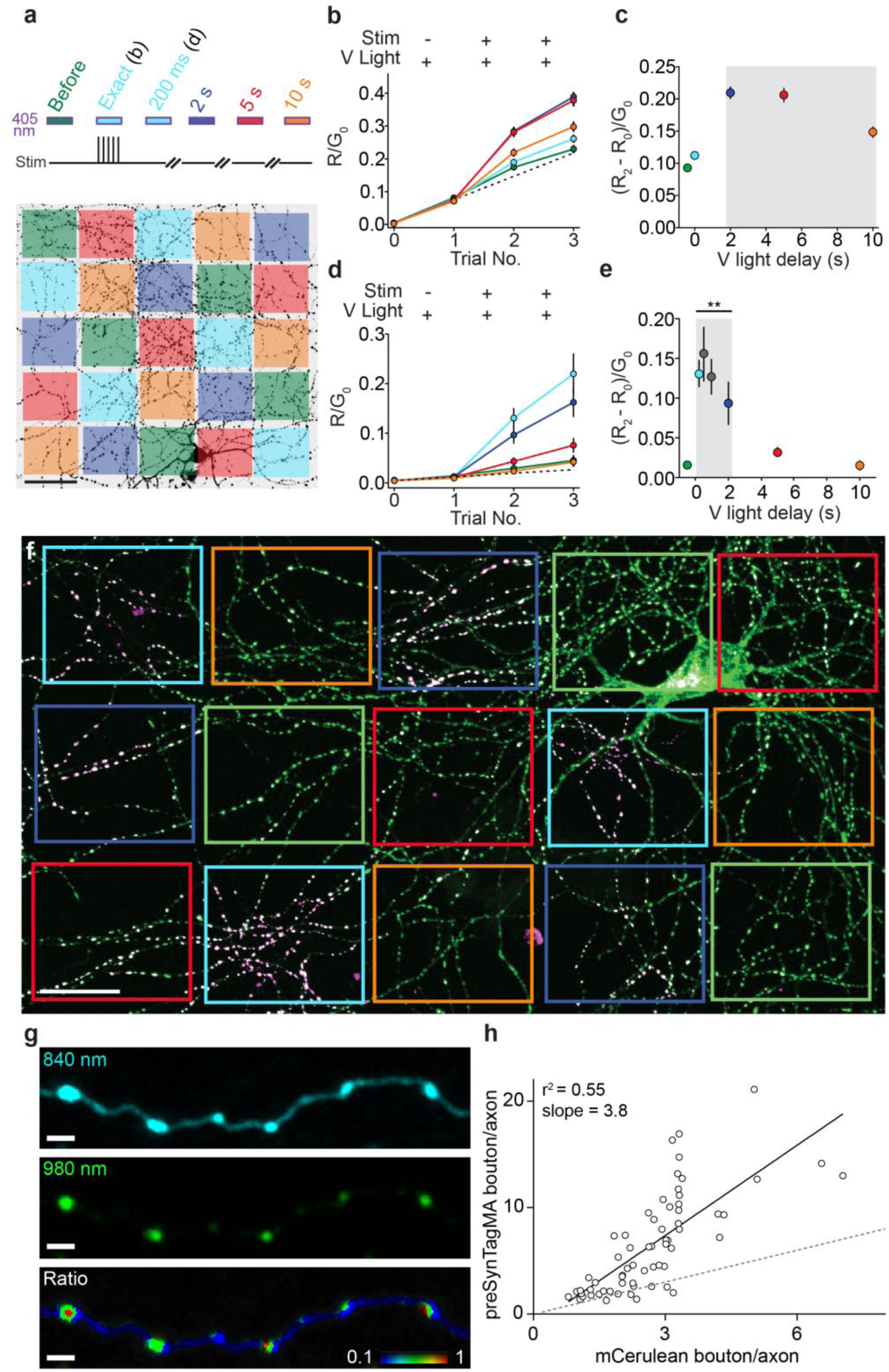
Temporal resolution of preSynTagMA photoconversion. **(a)** A spatial light modulator was used to illuminate parts of the axonal arbor (405 nm, 50 mW cm^-2^) at different times relative to a brief tetanic stimulation (5 APs). After each trial, new images were acquired. ‘Exact’ & ‘200 ms’ timing share the same color code (cyan) as we used ‘exact’ timing in sypCaMPARI experiments and a 200 ms delay in preSynTagMA experiments. **(b)** Ratio of red to green fluorescence (R/G_0_) from sypCaMPARI boutons illuminated at different times relative to the electrical stimulation. Trial 1 shows the effect of illumination without stimulation. Line color code as in **a**, n = 6 cells **(c)** Activity-dependent photoconversion (ΔR/G_0_) versus delay from start of stimulation to violet light from experiments in **b**. The grey box indicates the time window for efficient photoconversion of sypCaMPARI. **(d)** Neurons expressing preSynTagMA (synaptophysin-CaMPARI2 (F391W, L398V)) were stimulated as in **a-c**. Note the greatly reduced increase in R/G_0_ with violet light alone (Trial 1). n = 6 cells. **(e)** Activity-dependent photoconversion (ΔR/ G_0_) versus delay of preSynTagMA expressing neurons. n = 6-12 cells. **(f)** Cultured rat hippocampal neuron expressing preSynTagMA, trial # 3. Boxes indicate regions where photoconversion light was applied with different delays (color code as in **a**). **(g)** Two-photon image of hippocampal slice culture, showing axonal boutons in *stratum radiatum* originating from CA3 neurons expressing preSynTagMA and mCerulean as cytosolic filler. Upper panel: mCerulean (840 nm), middle panel: preSynTagMA (980 nm), lower panel: green/cyan ratio image, indicating labeled vesicle clusters. **(h)** The ratio of preSynTagMA bouton to axonal shaft fluorescence vs the ratio of mCerulean bouton to axon fluorescence for n = 64 boutons. Solid line: linear fit. Dashed line (slope = 1) indicates the expected location of boutons without synaptic vesicles. Data in **b-e** are presented as mean ± SE. ** p < 0.01 vs pre, Kruskal-Wallis ANOVA followed by Dunn’s multiple comparison. Scale bars: **(a)** 50 µm, **(f)** 25 µm, **(g)** 2 µm.

Parallel efforts to improve CaMPARI resulted in CaMPARI2, which contains a number of point mutations improving brightness, increasing kinetics, reducing activity-independent photoconversion and lowering the Ca^2+^ affinity^11^. We selected the variant CaMPARI2 (F391W_L398V) (*K*_d_ Ca^2+^ = 174 nM, *k*_on_ 66 s^-1^, *k*_off_ 0.37 s^-1^, photoconversion rate in slices 0.22 s^-1^ with Ca^2+^, 0.0021 s^-1^ without Ca^2+^)^11^ and fused it to synaptophysin to create preSynTagMA and expressed it in cultured hippocampal neurons. The temporal precision and dynamic range of preSynTagMA were both enhanced. PreSynTagMA showed no photoconversion in the absence of stimulation and the photoconversion time window was shortened to 0.2 – 2 s post-stimulation, corresponding to the faster dimming in response to 50 APs (**Fig. 2d, e, Supplementary Fig. 1h**). Using preSynTagMA, we could readily distinguish active from inactive axons in the presence of bicuculline (**Supplementary Fig. 1d-g**). Blocking action potentials (TTX) prevented SynTagMA photoconversion. Two hours after photoconversion, the R/G ratio was still 68% of the peak value, indicating photoconverted preSynTagMA marks activated boutons for several hours (**Supplementary Fig. 1i**). To quantify preSynTagMA localization in tissue, we co-expressed preSynTagMA together with cytosolic mCerulean in hippocampal slice cultures. Using two-photon excitation at 840 nm (mCerulean) and 980 nm (preSynTagMA green), we observed green fluorescent puncta at axonal boutons (**Fig. 2g**). Compared to mCerulean, preSynTagMA was 4-fold enriched in boutons vs. axonal shafts (**Fig. 2h**).

### Targeting SynTagMA to excitatory postsynapses

We chose to target SynTagMA to the postsynaptic protein PSD95, which has a higher retention time than most postsynaptic density proteins^16^. We decided against making a PSD95-CaMPARI fusion protein as overexpression of PSD95 is known to induce dramatic changes in function, size, and connectivity of dendritic spines^17^. Instead, we fused CaMPARI2 (F391W_L398V) (after deleting the nuclear export sequence) to a genetically encoded intrabody against PSD95 (PSD95.FingR)^15^. The resulting fusion protein, however, was not restricted to dendritic spines, where most excitatory synapses are located, but labeled the entire dendrite of CA1 pyramidal neurons (**Fig. 3a, d**). We reasoned that the lack of spine enrichment was due to a large fraction of unbound cytoplasmic protein. An elegant method to reduce cytoplasmic fluorescence is to fuse a zinc finger (ZF) and the transcription repressor KRAB(A) to the targeted protein and include a ZF binding sequence near the promoter^15,18^. It is presumed that the ZF-KRAB domains direct excess unbound cytosolic protein into the nucleus where the ZF binds to the ZF binding sequence and KRAB(A) suppresses transcription of the exogenous genes. We added these additional regulatory elements to the PSD95.FingR-CaMPARI2 (F391W_L398V) construct, which has no additional nuclear export or localization sequences added, and co-expressed it with mCerulean. Punctate green fluorescence was now observed predominantly in spines and nuclei were fluorescent as expected for a ZF-KRAB(A) containing protein. The ratio of spine-to-dendrite green fluorescence was about 4 times higher than mCerulean spine-to-dendrite ratios, suggesting CaMPARI2 was now localized to postsynapses (**Fig. 3b, e**). Serendipitously, we discovered that the upstream ZF binding sequence was dispensable for autoregulation (**Fig. 3c, f**), which simplified swapping of promotors. We named this minimal construct postSynTagMA and characterized it further. The important stabilizing function of autoregulation was confirmed in viral expression experiments (**Supplementary Fig. 2**). To test for potential effects of postSynTagMA on neuronal physiology, we measured passive and active electrical properties, miniature excitatory postsynaptic currents and spine densities of postSynTagMA/mCerulean-expressing neurons and neurons expressing only mCerulean (**Supplementary Fig. 3**). These control experiments, which were performed blind, yielded no significant differences between groups, indicating that expression of postSynTagMA did not alter neuronal physiology.

**Figure 3.**
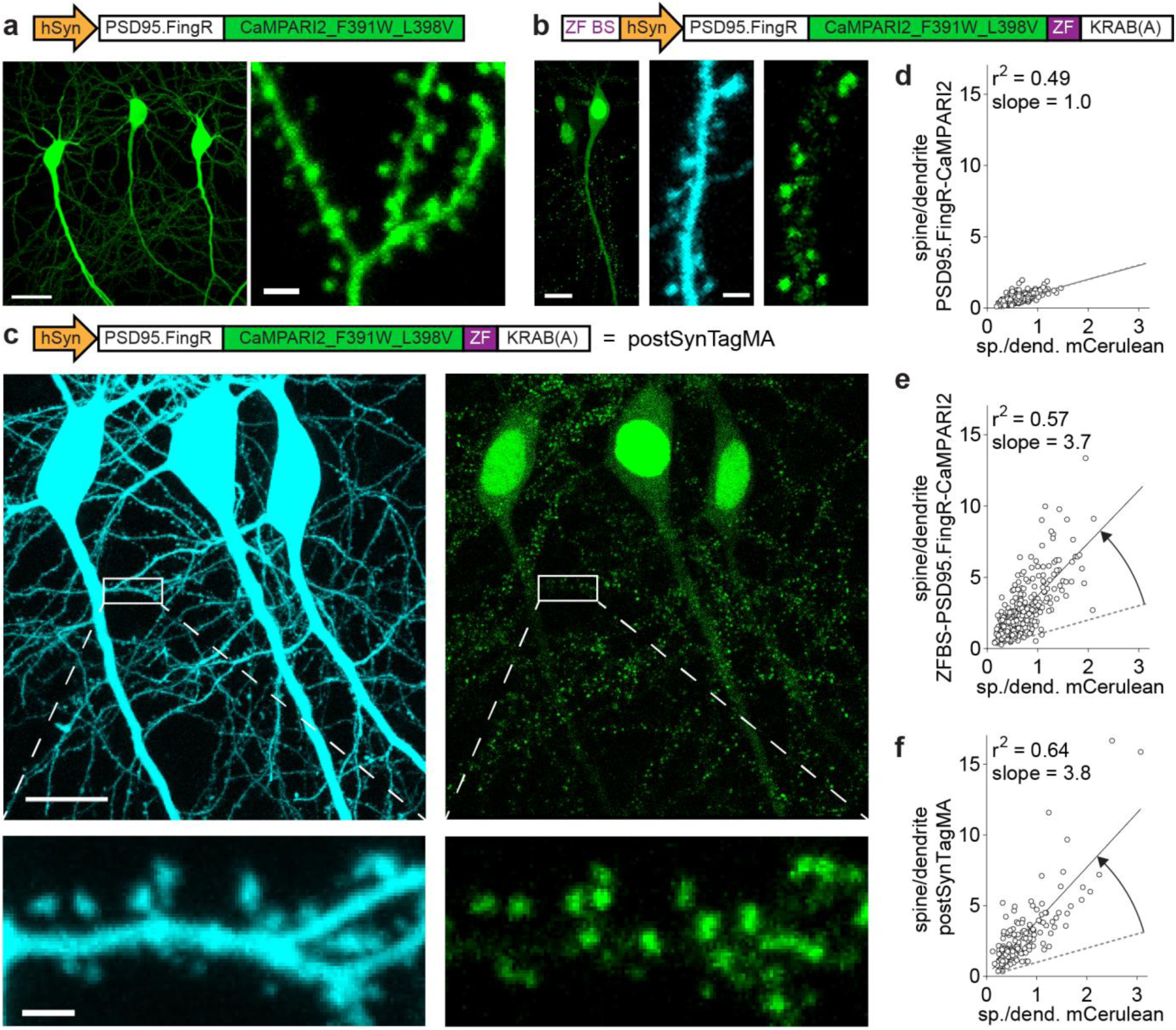
Postsynaptic targeting of SynTagMA using a PSD95 intrabody. **(a)** Two photon (2P) maximum intensity projection (980 nm) from CA1 neurons expressing a fusion protein of PSD95 fibronectin intrabody (PSD95.FingR) and CaMPARI2_F391W_L398V (without NES or epitope tags). Scale bars: 20 µm, 2 µm. **(b)** 2P maximum intensity projections from a CA1 neuron expressing PSD95.FingR-CaMPARI2 with a zinc finger binding sequence (ZF BS) added upstream of the promoter and a zinc finger (ZF) fused to a transcriptional repressor domain (KRAB(A)) and mCerulean (840 nm). Scale bar 12 µm (left panel) and 2 µm (center and right panels). **(c)** Postsynaptically targeted SynTagMA (postSynTagMA). As in **b**, with the ZF-KRAB(A) but no upstream ZF-BS. Note that postSynTagMA is still enriched in spines and the nucleus, leaving the cytoplasm almost free of SynTagMA. Scale bars: upper 20 µm, lower 2 µm. **(d)** The unregulated construct is expressed at high levels, leading to near-identical concentrations in dendrites and spines. For individual spines (circles), the spine-to-dendrite green fluorescence ratio was similar to the mCerulean spine-to-dendrite ratio. Line: linear fit. (**e**) The construct with autoregulatory elements shuts off its own production, resulting in strong enrichment in spines (3 neurons, 367 spines) (**f**) The construct without zinc finger binding sequence (postSynTagMA) is also auto-regulated, showing equally strong enrichment in spines (3 neurons, 179 spines).

### Using back-propagating action potentials to characterize postSynTagMA

We evoked backpropagating action potentials (bAPs) by brief somatic current injections to raise intracellular Ca^2+^, and applied 395 nm light pulses (500 ms) with a 1 s delay to characterize postSynTagMA photoconversion. In the absence of stimulation (0 bAPs), violet illumination did not change the R/G ratio (R_1_/G_1_ = R_0_/G_0_). Pairing violet light with bAP trains (15 repeats) lead to increased R_1_/G_1_ ratios (**Fig 4a**). To determine the best metric for quantifying SynTagMA photoconversion, we plotted several against the initial green fluorescence, which indicates synapse size (**Fig. 4b, Supplementary Fig. 4**). Considering only the change in red fluorescence (ΔR = R_1_ - R_0_), the apparent photoconversion positively correlated with synapse size and only large synapses in the 3 bAP and 50 bAP groups could be separated from the non-converted (0 bAP) synapses. The R_1_/G_1_ ratio, albeit showing better separation than ΔR, negatively correlated with size showing higher values and a higher chance to classify small synapses as photoconverted. When strong photoconversion occurs, small synapses with few indicator molecules may lose all green fluorescence, which is problematic when one wishes to divide by this measure. Evidence of this is seen in the extremely high values (i.e. 20 - 200) for R_1_/G_1_, especially in small to medium-sized synapses (**Fig. 4a**). As photoconversion both increases red and decreases green fluorescence, we reasoned that using all channels before and after conversion would provide an optimal metric and avoid dividing by values close to 0. Indeed, ΔR/(G_0_ + G_1_) was not correlated with PSD size. Using this measure, we detected a significant increase in photoconversion after 15 x 50 bAPs and even after 15 x 3 bAPs (**Fig. 4c and 4d**). To test whether violet light had adverse effects on cell health, we performed propidium iodide stainings of cultures exposed to different doses of violet light (**Supplementary Fig. 5**). The violet light dose used for SynTagMA photoconversion was well below the threshold for photodamage.

**Figure 4.**
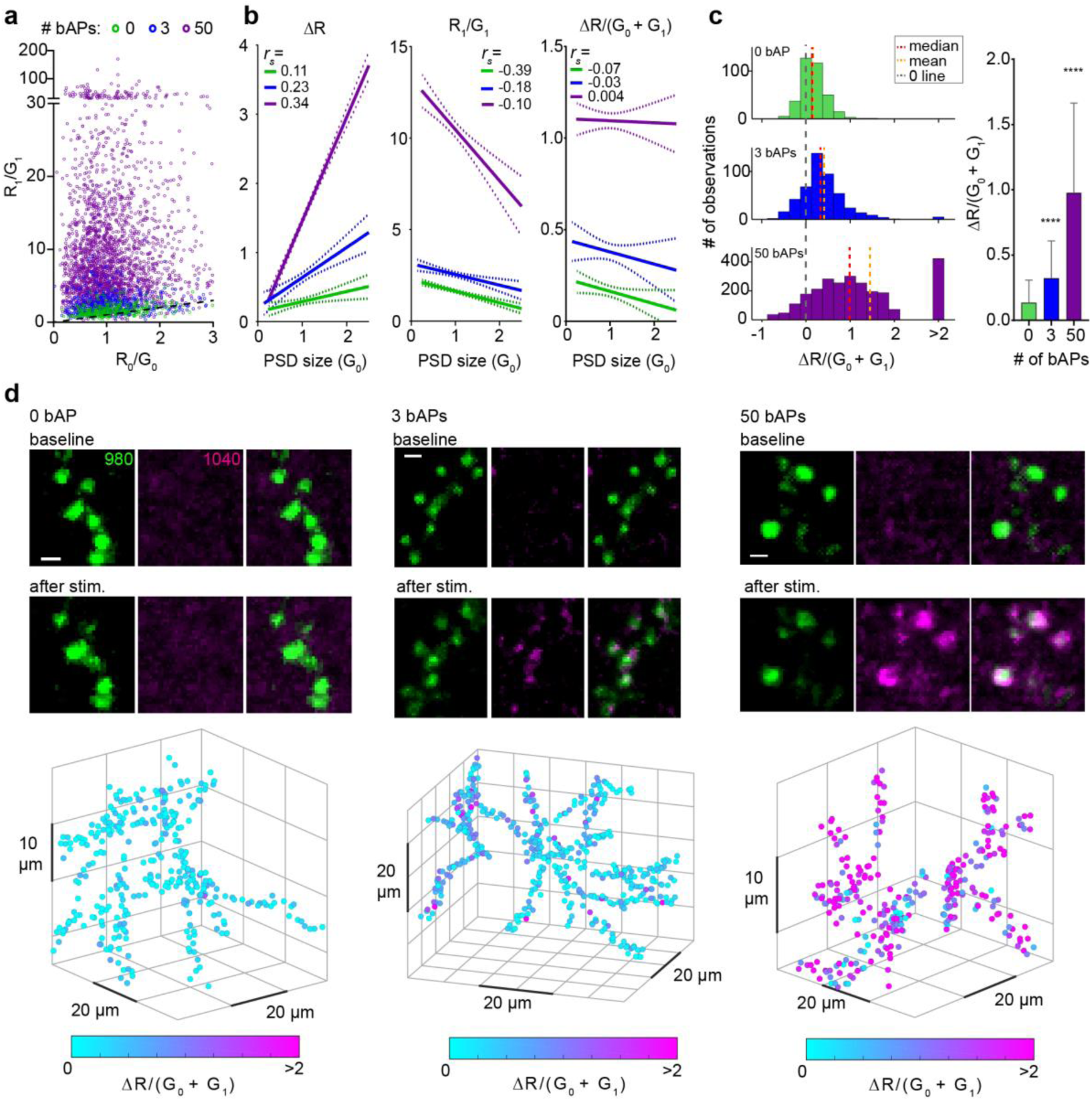
PostSynTagMA photoconversion with back-propagating action potentials (bAPs). 2P image stacks of postSynTagMA expressing CA1 pyramidal neurons were taken before and after 15 pairings of trains of bAPs with photoconverting violet light (395 nm, 16 mW mm^-2^, 500 ms duration, 1 s delay). Synaptic transmission was blocked. **(a)** R/G ratios of individual synapses before (R_0_/G_0_) vs after (R_1_/G_1_) photoconversion. Note the variability in photoconversion within conditions. Dotted black line is the unity line. **(b)** Three different metrics vs PSD size (Magenta: 50 bAPs. Blue: 3 bAPs. Green: 0 bAP; linear regression lines with 95% confidence intervals). ΔR is positively correlated with PSD size (0 bAPs: *r*_*s*_ = 0.11, p = 0.03; 3 bAPs: *r*_*s*_ = 0.23, p < 0.0001; 50 bAPs: *r*_*s*_ = 0.34, p < 0.0001). Normalizing ΔR by (G_0_ + G_1_) removes the correlation with PSD size (0 bAP: *r*_*s*_ = −0.07, p = 0.164; 3 bAPs: *r*_*s*_ = −0.03, p = 0.583; 50 bAPs: *r*_*s*_ = 0.004 p = 0.876). *r*_*s*_ is the correlation coefficient (Spearman’s rho). **(c)** Distributions of photoconversion (ΔR/(G_0_+G_1_) for each condition (0 bAP: median = 0.139, mean = 0.164; 3 bAPs: median = 0.327, mean = 0.406; 50 bAPs: median = 0.984, mean = 1.452) and ΔR/(G_0_+G_1_) vs the number of bAPs (data are median and interquartile range; ****p < 0.0001, Kruskal-Wallis vs. 0 bAP). 0 bAP: n = 1 cell, 356 synapses; 3 bAPs: n = 1 cell, 472 synapses; 50 bAPs: n = 3 cells, 2587 synapses. **(d)** Example green, red and merged images of SynTagMA labelled spines photoconverted with 0, 3 or 50 bAPs. Scale bars: 1 µm. Below: Photoconversion of individual spines was color-coded and plotted at their 3D-position in the dendritic arbor.

**Figure 5.**
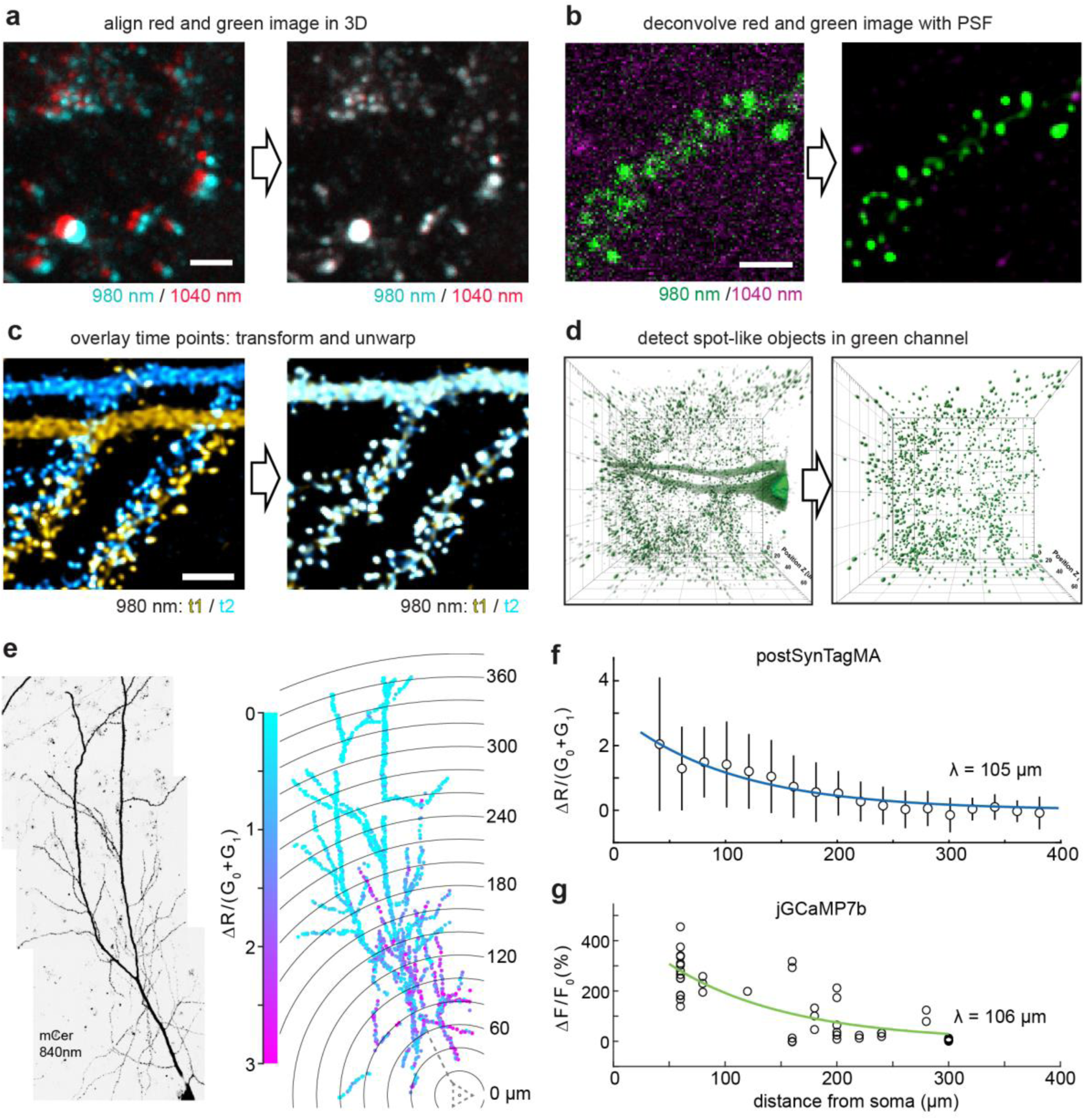
Analysis workflow for automated detection and analysis of SynTagMA photoconversion. **(a)** Green and red fluorescence is collected in separate images, switching between two Ti/Sapph lasers (980 nm / 1040 nm). Images are registered in 3D to correct for chromatic aberration and laser alignment. Scale bar: 4 µm. **(b)** Median filtering and deconvolution is then applied to all images (both green and red channels). Scale bar: 3 µm. **(c)** To superimpose multiple time points in 3D, translation, rotation and unwarping are applied. Scale bar 4 µm. **(d)** Synapses (ROIs) are detected as spherical objects, i.e. ‘spots’ from which fluorescence values are extracted and analyzed. **(e)** Maximum intensity projection of a CA1 pyramidal cell expressing cytosolic mCerulean (inverted gray scale) and postSynTagMA. The cell was stimulated with 50 bAPs at 100 Hz and illuminated with 395 nm (as in Fig. 4). For each identified synapse, ΔR/(G_0_ + G_1_) was analyzed, color-coded, and plotted at its original location. Distance from soma is indicated as concentric rings. **(f)** Photoconversion decreased exponentially from the soma with a distance constant of λ = 105 µm (median ± interquartile range, n = 1860 synapses, R^2^ = 0.91). **(g)** Spine calcium transient amplitudes during bAP trains (jGCaMP7b) decreased exponentially with distance from the soma with λ = 106 µm, (n = 55 synapses, R^2^ = 0.66).

### Analysis workflow for automatic synapse detection and quantification of SynTagMA

While the efficient targeting of SynTagMA allows simultaneous interrogation of a large population of synapses, it also presents an analysis challenge. To place regions of interest (ROIs) on individual synapses (**Fig. 1 and 2**), simple spot detection algorithms (e.g. Imaris, ImageJ) can be used, but matching of objects across several time points is not straightforward. When the number of synapses reaches into the thousands, a manually curated approach is no longer feasible. We developed an image analysis workflow to tackle this problem. Images acquired using two-photon excitation at different wavelengths must first undergo correction for laser alignment and chromatic aberration (**Fig. 5a**). As synapses in tissue are quite motile even on short time scales^19,20^, a non-rigid 3D transformation, termed ‘unwarping’, was applied to re-align and preserve synapse identity across time points. To achieve the highest quality transformation, images were first deconvolved and then underwent ‘unwarping’ (**Fig. 5b-c**). Synapses were then detected on transformed datasets using the Spot feature of Imaris (Oxford Instruments) (**Fig. 5d**). Although each of these steps can be performed using a combination of freely and commercially available software packages, it is a time-consuming process that generates several gigabytes of data in intermediate steps. We therefore developed SynapseLocator (available at GitHub), a Matlab-based program to streamline the aforementioned steps using freely available ancillary software tools (Fiji^21^, DeconvolutionLab2^22^, FeatureJ, elastix^23^). A machine-learning approach^24^ was implemented to generate synapse templates (boutons or postsynaptic sites) that were used to automatically detect and extract fluorescence values. Transformed images and fluorescence values from identified synapses (ROIs) were saved for statistical analysis and imported into ImageJ or Imaris for 3D visualization (**Fig. 5d**). A specific difficulty for automated analysis is the presence of spot-like red autofluorescence at 1040 nm excitation. Synapses with elevated R_0_/G_0_ values at baseline were considered too close to autofluorescent objects and were automatically excluded from further analysis. Automation of the synapse identification process greatly reduced the analysis time (from days for ∼500 hand-curated ROIs at several time points to minutes for ∼4000 automatically detected ROIs from the same dataset, using a personal computer). Manual and automated analysis produced comparable results (**Supplementary Fig. 6**).

**Figure 6.**
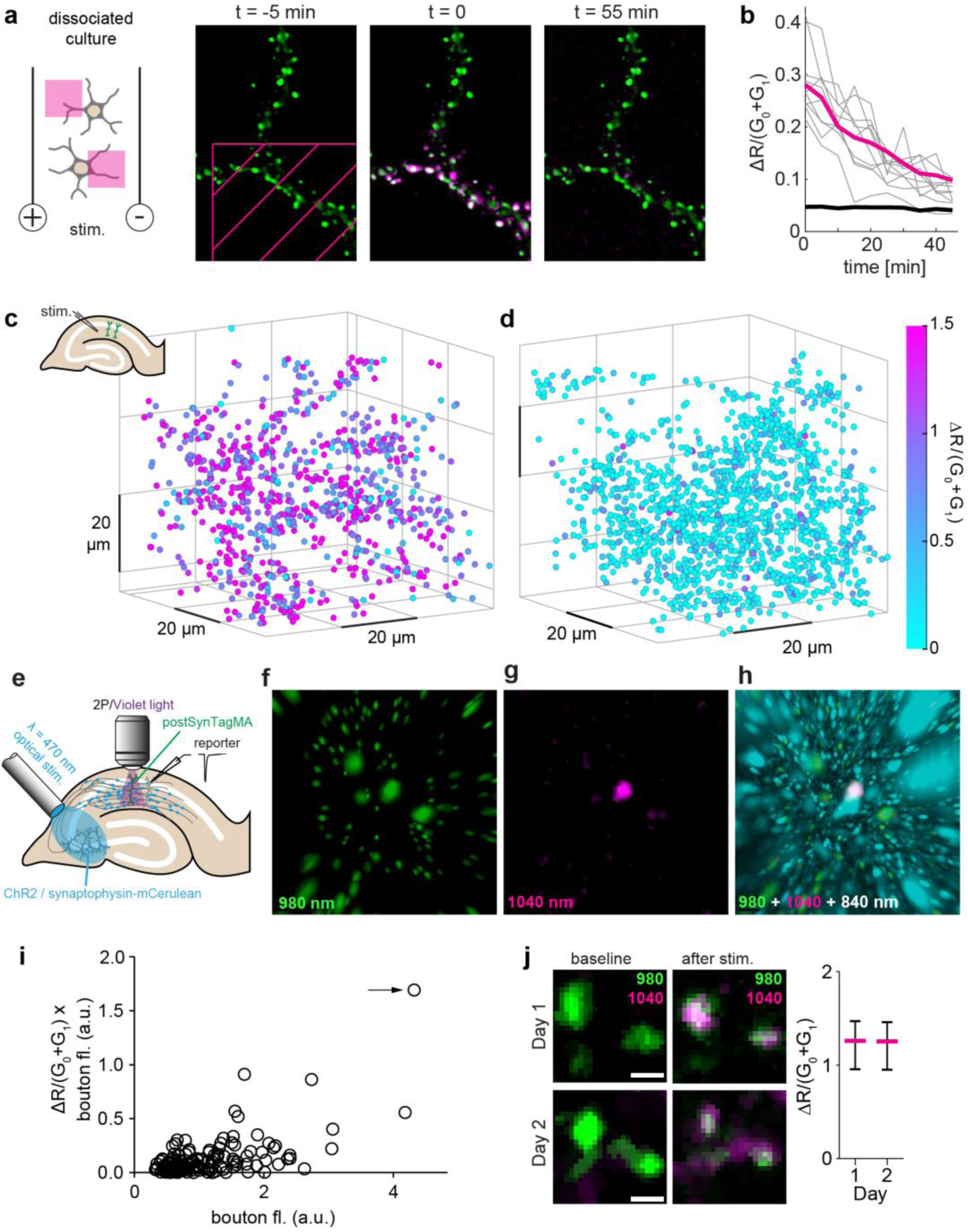
Using postSynTagMA to map active synapses. **(a)** Time series of postSynTagMA spots on the dendrite of a cultured neuron. At t = 0, one violet light pulse was applied via DMD inside the hatched area (405 nm, 18.6 mW mm^-2^, 0.5 s, t = 0 min) while stimulating the culture with a single train of action potentials delivered at 50 Hz for one second. **(b)** Inside the illuminated area (magenta line), red fluorescence generated by photoconversion of postSynTagMA decayed exponentially with τ = 29.4 min (R^2^ = 0.99, n = 138, 2 experiments). Gray lines show individual examples (n = 1). For objects outside the illuminated square, ΔR/(G_0_+G_1_) was constant (black line, n = 712, 2 experiments). **(c)** Photoconversion after Schaffer collateral stimulation in organotypic slice culture. Color-coded ΔR/(G_0_ + G_1_) of spines plotted at their location in the 65 x 65 x 78 µm volume of tissue imaged. Strong stimulation of synaptic inputs in *stratum radiatum* was paired with 395 nm light (100 ms, delay 1s, 50x). Note that ΔR/(G_0_ + G_1_) is equal or greater than 1.5 (magenta) for most of the synapses (n = 897). **(d)** As in **c**, but with weak extracellular stimulation, below the threshold for postsynaptic APs (n = 1502). **(e)** CA3 neurons expressing ChR2-ET/TC and synaptophysin-mCerulean were stimulated with 470 nm flashes to evoke subthreshold (∼500 pA) excitatory synaptic responses in a patch-clamped CA1 neuron (reporter). Simultaneously, an adjacent postSyntagMA-expressing CA1 neuron was illuminated with violet light (50 repeats, 100 ms, 395 nm, 16 mW mm^-2^) through the objective. **(f)** Green spots (G_0_+G_1_) segmented with Imaris™. **(g)** Red fluorescence after photoconversion (R1), masked by the green channel. A strongly photoconverted synapse (spot #1) is seen in the center. **(h)** Spot #1 (magenta) is in close contact with a presynaptic ChR2-expressing terminal (cyan). Non-converted (green) spots are not in direct contact with cyan terminals. **(i)** Quantitative analysis of panels f-h (n = 167 spots). Arrow indicates spot #1. **(j)** Red/green overlay before and after photoconversion of the same stretch of dendrite on successive days. Strong extracellular synaptic stimulation was paired with 395 nm light pulses (500 ms, 1s delay, 15x). Photoconversion of postSynTagMA was near-identical on day 1 and day 2 (median ± inter-quartile range, n = 37 synapses). Scale bar: 1 µm.

### Back-propagating action potentials increase Ca^2+^ at proximal synapses

For a given number of bAPs, the amount of photoconversion was quite variable between individual spines (**Fig. 4a**), likely reflecting spine Ca^2+^ transients of different amplitude and duration. We also observed that whereas synapses close to the soma were strongly photoconverted, conversion decreased with distance, suggesting that bAP bursts do not increase [Ca^2+^] in distal branches of the apical dendrite (**Fig. 5e and 5f**). To test whether SynTagMA conversion is a linear reporter of local Ca^2+^ signal amplitude, we performed equivalent experiments on CA1 neurons expressing jGCaMP7b. Indeed, Ca^2+^ imaging during trains of bAPs yielded the same exponential decay of amplitude as a function of distance from the soma, validating our observation that bursts of bAPs do not invade distal dendrites (**Fig. 5g**). In contrast to imaging Ca^2+^ synapse-by-synapse, postSynTagMA photoconversion reliably reports the amplitude of bAP-induced Ca^2+^ elevations across the entire dendritic tree, resulting in higher coefficients of determination (R^2^) in the statistical analysis (R^2^ = 0.91 vs 0.66).

### Using postSynTagMA to map synaptic activity

Having established the linear Ca^2+^-dependence of photoconversion in patch-clamped CA1 neurons, we wanted to assess how long converted postSynTagMA would persist in individual spines. We illuminated rectangular areas of cultured neurons while stimulating the culture at 50 Hz, limiting postSynTagMA photoconversion to the illuminated dendritic sections (**Fig. 6a**). We observed that ΔR/(G_0_ + G_t_) returned to baseline with a time constant of 29 min (**Fig. 6b**), in contrast to the much slower turnover of preSynTagMA (**Supplementary Fig. 1i**) and of photoconverted soluble CaMPARI^11^. This relatively fast decay of the spine signal, which is consistent with the short retention time of synaptic PSD95 in the neocortex of young mice^25^, limits the post-photoconversion acquisition time to about 30 minutes.

We next tested whether presynaptic stimulation would raise spine Ca^2+^ sufficiently to trigger postSynTagMA photoconversion. In hippocampal slice culture, we activated Schaffer collateral axons and illuminated the dendrite of postSynTagMA-expressing CA1 pyramidal cells (100 ms violet light after 1 s delay, 50 repeats). Strong synaptic stimulation resulted in widespread photoconversion in dendritic spines (**Fig. 6c**). However, as postsynaptic neurons may have been driven to spike in these experiments, we could not distinguish spines that received direct synaptic input from spines that were passively flooded with Ca^2+^. Weak stimulation resulted in a very sparse and distributed conversion pattern, suggesting active synapses (**Fig. 6d**). In the example, a 65 x 65 x 78 µm^3^ volume contained 1500 postSynTagMA-labeled synapses, fourteen of which showed values above 3s (ΔR/(G_0_ + G_1_) = 0.97) and were therefore classified as active. Given the relatively low release probability of Schaffer collateral synapses, only a third of this 1% of activated synapses will release transmitter at any given stimulation pulse. This low number of synchronized inputs is not expected to generate APs or dendritic spikes in the postsynaptic neuron, which is consistent with the distributed pattern of photoconverted synapses.

To demonstrate that postSynTagMA indeed labels active synapses, it would be desirable to have an independent marker of synaptic activation. We turned to optogenetic stimulation to allow visualization of active presynaptic terminals. CA3 neurons expressing channelrhodopsin2 ET/TC (ChR2) and synaptophysin-mCerulean^26^ were stimulated by blue light pulses (**Fig. 6e**). Light stimulation intensity was adjusted to be below the threshold for AP generation in a patch-clamped CA1 pyramidal cell (‘reporter’ neuron). Combining light stimulation of CA3 neurons with violet light illumination of postSynTagMA-expressing CA1 pyramidal cells resulted in very sparse photoconversion of Schaffer collateral synapses (**Fig. 6f, g**). Next to strongly photoconverted spines, cyan fluorescent boutons were observed, suggesting that these synapses were directly innervated and activated by ChR2-expressing presynaptic CA3 neurons (**Fig. 6h**). Non-photoconverted spines were also distant from activated terminals and thus represented ‘true negatives’ (**Fig. 6i, Supplementary Movie 1**). As red postSynTagMA has a relatively rapid turnover (**Fig. 7b**), it should be possible to generate multiple maps of active synapses over time. Indeed, using electrical stimulation of Schaffer collateral axons, we were able to convert (median ΔR/(G_0_ + G_1_) = 1.26) and, 18 hours later, reconvert (median ΔR/(G_2_ + G_3_) = 1.25) SynTagMA-labeled spines on CA1 pyramidal cells (**Fig. 6j**). The remarkably similar degree of conversion on consecutive days suggests the possibility of repeated activity mapping with postSynTagMA.

**Figure 7.**
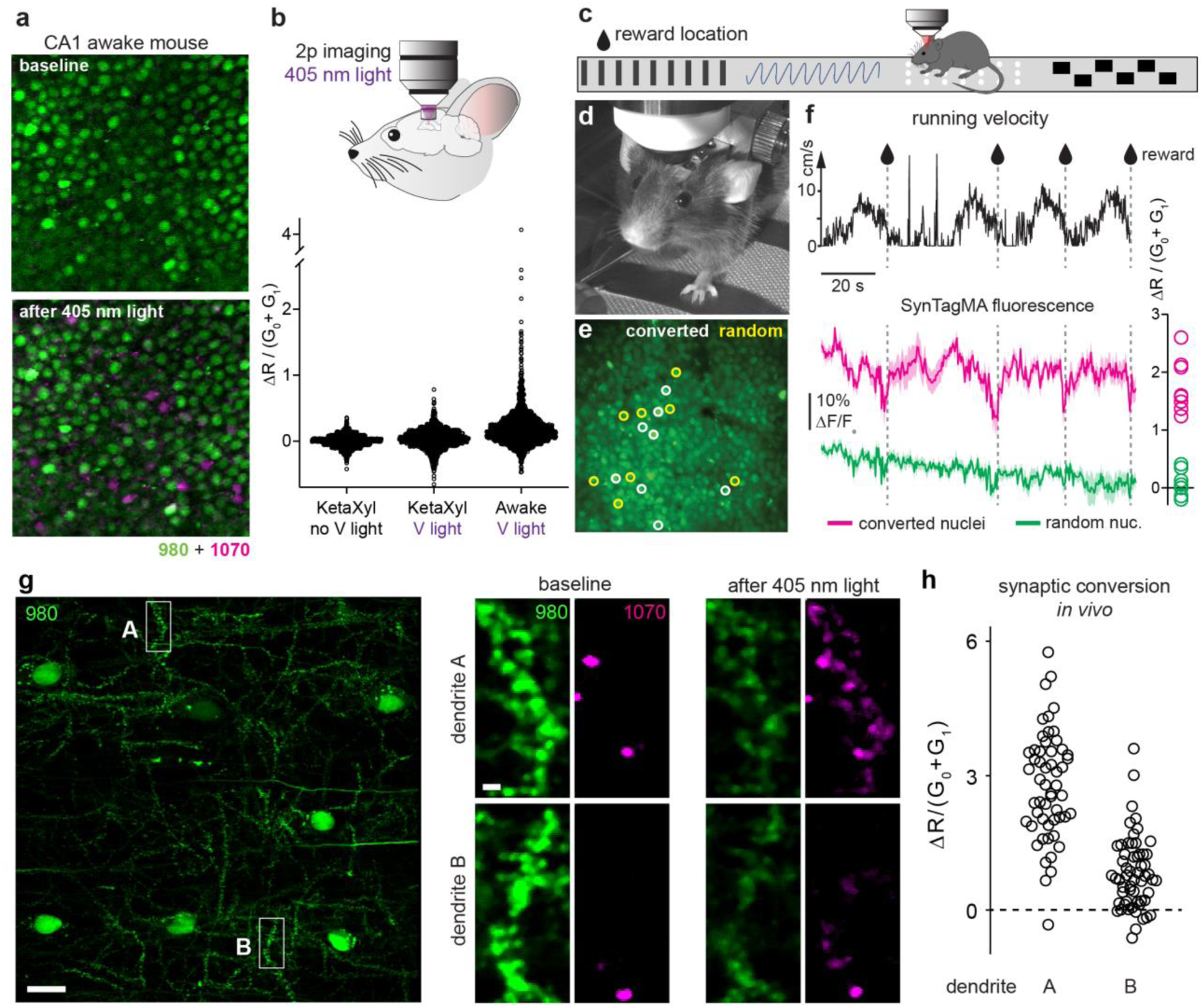
SynTagMA identifies active neurons during behavior. **(a)** Nuclei of postSynTagMA expressing CA1 neurons imaged *in vivo* through a chronic cranial window using 980 nm and 1070 nm to excite green and red SynTagMA fluorescence, respectively. Ten 2 s, 405 nm, 12.1 mW mm^-2^ violet light pulses were applied in an awake mouse after which a small percentage of nuclei became photoconverted (magenta). **(b)** Quantification of photoconversion relative to baseline under ketamine-xylazine anesthesia (Keta/Xyl no V light), after 405 nm light as in A (V light) and when the mouse was awake. **(c)** Closed loop paradigm: A head-fixed mouse was trained to stop at a certain position on the running belt to receive a water reward (teardrop). Nuclear fluorescence in CA1 was continuously monitored during 4 laps, followed by 15 laps where 405 nm, 12.1 mW mm^-2^, 2 s light pulses were triggered together with the water reward. **(d)** Mouse engaged in the task. Note spout for water delivery. **(e)** Two-photon image of CA1 cell body layer during running. Eight nuclei that later became photoconverted are marked by white circles. Yellow circles are the eight randomly selected non-converted nuclei used for comparison of calcium signals. **(f)** Running velocity, corresponding average calcium responses (SynTagMA green fluorescence) of laps 1-4 and subsequent photoconversion of the corresponding eight converted and eight non-converted CA1 nuclei indicated in E. Black trace is running speed during the first 4 laps with times of reward delivery (teardrops/dashed lines). Magenta and green traces are the average green SynTagMA fluorescence of the 8 photoconverted nuclei and the 8 non-converted nuclei, respectively. At right is the photoconversion of the individual nuclei. Note the consistent dips in the magenta trace (i.e. high calcium) just before the water reward/photoconversion light would be triggered. **(g)** PostSynTagMA-expressing interneurons imaged through a chronic cranial window under isoflurane anesthesia. At high magnification, green fluorescence reveals PSD spots on the dendrite. Red spots at baseline are autofluorescent material, not related to SynTagMA. Violet light (20 flashes, 0.2 Hz, 3 s duration, 0.42 mW mm^-2^) was applied to photoconvert postSynTagMA. Scale bars: 20 µm and 2 µm, respectively. **(h)** Photoconversion of synaptic spots on dendrite A (n = 54) and B (n = 58) indicate higher activity levels in dendrite A.

### PostSynTagMA maps active neurons and synapses *in vivo*

The nuclear sequestration of postSynTagMA prompted us to test whether it could be used to identify active neurons *in vivo*. When a head-fixed mouse running on a linear treadmill was illuminated with 405 nm light, a small percentage of the CA1 neuronal nuclei expressing postSynTagMA turned red (**Fig. 7a**). In the same mouse, there was no photoconversion during ketamine-xylazine anesthesia (**Fig. 7b**), consistent with the strongly reduced activity levels (**Supplementary Movie 2**). We next trained mice to stop and receive a water reward at a particular location (**Fig. 7c-d**). After reaching criterion, we monitored green fluorescence continuously during four laps (**Fig. 7e and 7f**), followed by 15 laps with reward-triggered light. The neurons that were photoconverted also showed dimming (i.e. increased Ca^2+^) immediately prior to each reward (**Fig. 7f**, magenta) whereas a matched number of randomly selected non-converted neurons did not (**Fig. 7f**, green). Therefore, in awake behaving animals, postSynTagMA photoconversion selectively labels neurons with high calcium transients (**Supplementary Movie 3**). In contrast to the analysis of cytoplasmic calcium indicators, we found that automatic segmentation and analysis of the nuclear SynTagMA was trivial.

The density of neuropil labeling in these mice prompted us to switch to expressing postSynTagMA in interneurons (**Supplementary Fig. 7**) to test whether postSynTagMA could be used to identify individual active synapses/dendrites in sparsely labeled neurons in vivo. We photoconverted sparsely labeled interneurons in *stratum oriens* interneurons under isoflurane anesthesia which retains neuronal activity at close to awake levels but reduces motion artifacts (**Fig. 7g-h, Supplementary Movie 2**). Photoconversion of clusters of synapses was observed, which would be consistent with local dendritic calcium events that have been described in interneurons in vivo^27^. Thus, under conditions of sparse SynTagMA expression, it is possible to resolve individual photoconverted synapses *in vivo*. During photoconversion, movement of the brain is not problematic and SynTagMA can be used to tag active neurons and synapses during behavior. To later resolve the SynTagMA label at the level of individual synapses, minimizing tissue movements with anesthesia may improve data quality.

## DISCUSSION

To create a synaptically localized, photoconvertible calcium sensor, we fused CaMPARI2 to the vesicle protein synaptophysin (preSynTagMA) for presynaptic targeting or to a nanobody against the postsynaptic scaffold of excitatory synapses (postSynTagMA). Both targeting strategies have previously been used to label synapses with fluorescent proteins^15,28^, and we verified that the autoregulatory elements of postSynTagMA do not cause widespread misregulation of endogenous gene expression and that SynTagMA-expressing neurons are physiologically normal. When SynTagMA is combined with a second fluorescent protein, here mCerulean, postSynTagMA puncta can be assigned to particular dendrites or neurons. PreSynTagMA should prove useful for identifying active axons in tissue and may also prove useful for distinguishing high from low release probability boutons based on the graded photoconversion. PostSynTagMA should likewise facilitate identification and mapping of active synapses, but on the postsynaptic side. Antibodies specific to the photoconverted red form will also enable further ultrastructural analysis of the marked synapses^11^.

Identifying active synapses is a long-standing quest in neuroscience and many approaches have been put forward. In two-dimensional neuronal cultures, it is relatively straightforward to monitor the activity of a large number of synapses with a high speed camera, e.g. by employing genetically encoded sensors of vesicular pH^29,30^. In intact brain tissue, however, monitoring more than one synapse with high temporal resolution becomes very challenging^6,31^. To create a more permanent tag of active synapses, Ca^2+^ precipitation in active spines was used to create an electron-dense label that can be detected by electron microscopy ^32^. As this method requires tissue fixation, it does not allow monitoring changes in synaptic activity over time. Expressing the two halves of split-GFP on the extracellular surface of pre- and postsynaptic neurons (mGRASP) labels contact points between the neurons as green fluorescent puncta^33^. Refined transsynaptic labeling methods with multiple colors have led to remarkable insights about network connectivity^34,35^. Whereas mGRASP and eGRASP report anatomical proximity, SynTagMA photoconversion is Ca^2+^-dependent and therefore sensitive to synaptic activity.

One of the many interesting applications of postSynTagMA will be to create extensively detailed input maps of individual neurons at single synapse resolution. The question of whether inputs carrying similar information segregate to branches of the dendritic tree is currently being investigated using spine-by-spine Ca^2+^ imaging^6,36,37^. By freezing the activity status of all spines during a particular labelling protocol, SynTagMA may simplify such experiments by reading out fluorescence ratios in thousands of spines in a single high-resolution stack. Importantly, postSynTagMA maps can be repeated, opening the possibility to study the functional dynamics of excitatory connections rather than just morphology and turnover of dendritic spines. In addition to synapses on mushroom-shaped spines, postSynTagMA provides access to synapses on structures that are difficult to probe with Ca^2+^ dyes or diffusible GECIs, such as stubby spines or shaft synapses^38^. Thus, postSynTagMA may open the possibility to study the long-term dynamics of interneuron excitation, a key homeostatic mechanism that involves Ca^2+^-induced-Ca^2+^ release from intracellular stores^39^. In addition to Ca^2+^ influx though voltage- or ligand-gated channels, Ca^2+^ release is thought to be important in both presynaptic^40^ and postsynaptic compartments^41–43^, and SynTagMA may help investigating the sources of Ca^2+^ in individual synapses.

To test the linearity of calcium-dependent photoconversion, we used bursts of action potentials (APs) which are known to be strongly attenuated on their way into the dendrite^44,45^. Indeed, postSynTagMA photoconversion was greatest in spines relatively close to the soma while distal synapses were not converted at all. The distance constant estimated from a Sholl analysis of postSynTagMA conversion (λ = 105 µm, **Fig. 5f**) was the same as that determined by calcium imaging of individual spines (λ = 106 µm, **Fig. 5g**). Previous studies^44,45^ have measured back-propagation along the apical dendrite, but not along thin oblique dendrites or into individual spines. Studies with single-spine resolution^46,47^ have not included large-scale spatial information, as only few synapses per cell can be investigated. In future SynTagMA studies, reconstructions of the neuronal morphology in 3D would allow accurate measurements of the dendritic path length to every synapse and combine that data with functional information. We observed that even close to the soma, conversion of spines was not very uniform and the same was true for boutons. Understanding the sources of this heterogeneity, and whether these correspond to particular classes of synapses, may lead to important insights into synapse-specific calcium regulation.

An unexpected application arose from the nuclear localization of postSynTagMA. As we show, nuclear SynTagMA conversion can be used to visualize the location of the most active neurons in behaving animals. As the cytoplasm is almost free of label, automatic segmentation is very easy. Nuclear calcium elevations may be of particular relevance during learning, as they are driven by burst firing and trigger transcription of plasticity-related genes^48^. There are already several strategies for labeling active neurons^49^, but their time resolution is much worse than the 2 s photoconversion time window of SynTagMA.

### Limitations of the method

There are some caveats to working with SynTagMA. Before or during imaging, it is important to photoswitch the protein to the bright state using low intensity violet light^10,11^. The quite rapid turnover of postSynTagMA means that after photoconversion, images should be collected within about 30 minutes. SynTagMA analysis is complicated by endogenous red fluorescence which can be mistaken for photoconverted SynTagMA. For this reason, we found acquisition of a pre-photoconversion stack essential for reliable analysis: Synaptic ROIs (detected in the green channel) that already show elevated red fluorescence *before* stimulation (large R_0_/G_0_) can be classified as ‘contaminated’ and excluded from further analysis (see **Fig. 7g** for example red “spots” in both pre- and post-photoconversion images). If for experimental reasons only a single time point can be acquired, we recommend excluding objects that have high red fluorescence in the voxels surrounding green puncta. When acquiring two-photon image stacks, we collected each optical section twice, exciting either the green or the red form of SynTagMA by rapidly switching the power of two different Ti:Sapph lasers. Near-simultaneous acquisition of red and green fluorescence prevented movement artefacts and made it straightforward to correct for chromatic aberration. When the two color channels are collected in successive stacks, Ca^2+^-dependent dimming driven by spontaneous activity may affect one and not the other channel. Particularly when imaging in vivo, the time required to re-tune a single laser to the different excitation wavelengths might make analysis of individual synapses impossible.

If the postsynaptic neuron fires APs during strong (supra-threshold) synaptic activation, the resulting global calcium elevation could interfere with SynTagMA-based input mapping at proximal synapses. As we show (**Fig. 5e**), this is much less of a problem in the distal dendritic tree. For input-mapping experiments, it is advisable to repeat weak, subthreshold stimuli many times, each time paired with a (delayed) pulse of violet light. To avoid phototoxicity, it is important to keep track of the total light dose applied during each protocol, using shorter pulses (or lower light intensities) for protocols with many repetitions. SynTagMA is amenable for viral delivery using recombinant AAVs. Using different promoters, we demonstrate pan-neuronal expression for network analysis (**Fig. 7a-f**) and sparse expression of postSynTagMA in hippocampal interneurons (**Fig. 7g, Supplementary Fig. 7**). For sparse expression in pyramidal cells, dual-AAV labeling systems could be employed^50^.

## AUTHOR CONTRIBUTIONS

**Conceptualization**, TGO, CEG, MH and SW; **Methodology**; **Software** CS, IAG; **Investigation**, APA, BCF, RJO, WY, SFU, PLM, BM, MAM, LCP, JSW, MBH; **Resources** ERS; **Writing – Original Draft**, CEG, TGO, MBH, JSW, BCF, APA, CS, SFU; **Writing – Review & Editing**, APA, BCF, RJO, WY, IAC, SHU, PLM, BM, MAM, LCP, CS, ERS, JSW, CEG, MBH, TGO; **Visualization**, BCF, APA, CS, TGO, MBH; **Supervision** ERS, JSW, CEG, MBH, TGO; **Project Administration** CEG; **Funding Acquisition**, ERS, JSW, CEG, MBH, TGO.

## ACKNOWLEDGEMENTS

We thank Iris Ohmert, Sabine Graf and Kathrin Sauter for excellent technical assistance. Dr. Ingke Braren of the UKE Vector Facility produced AAV vectors. mCerulean was kindly provided by Dr. Wolfgang Wagner at the ZMNH. PSD95.FingR was kindly provided by Dr. Don Arnold at USC. Synaptophysin-GCaMP3 was kindly provided by Loren Looger at Janelia Farm. Funding was received from: The Deutsche Forschungsgemeinschaft (DFG) grants SPP 1665 220176618, SPP1926 273915538, SFB 936 178316478, SFB 1328 335447717, FOR 2419 278170285; the European Research Council (ERC-2016-stG 714762); the National Institute of Health (GM13132); the Howard Hughes Medical Institute. BCF was supported by an individual fellowship from the DAAD (Research Grants - Doctoral Programmes in Germany, 57214224). RJO was supported by the Esther and Joseph Klingenstein-Simons Foundation (FP00003669) and U.S. Department of Education GAANN grant (P200A150059). LCP was supported by NSF IOS grant 1750199. BM holds a postdoctoral fellowship from the Research Foundation-Flanders (FWO Vlaanderen)

## METHODS

### Experimental models

#### Dissociated rat hippocampal cell cultures

Neurons from P1 Sprague-Dawley rats of either sex were isolated from hippocampal CA1-CA3 regions with dentate gyrus removed, dissociated (bovine pancreas trypsin; 5 min at room temperature), and plated on polyornithine-coated coverslips inside a 6 mm diameter cloning cylinder. Calcium phosphate-mediated gene transfer was used to transfect 5-7 day old cultures. All measurements unless otherwise noted, are from mature 13-21 day old neurons. Procedures with Sprague-Dawley rats were approved by Dartmouth College’s Institutional Animal Care and Use Committee (IACUC). Cells were maintained in culture media consisting of Earle’s MEM (Thermofisher 51200038), 0.6% glucose, 0.1 g l^-1^ bovine transferrin, 0.25 g l^-1^ insulin, 0.3 g l^-1^ glutamine, 5% fetal calf serum (Atlanta Biologicals), 2% B-27 (Life Technologies), and 4 μM cytosine β-D-arabinofuranoside added 48 hours after plating in 6 mm diameter cloning cylinders (Ace Glass).

#### Rat hippocampal slice cultures

Hippocampal slice cultures from Wistar rats of either sex were prepared at postnatal day 4–7 as described (Gee et al., 2017). Briefly, rats were anesthetized with 80% CO_2_ 20% O_2_ and decapitated. Hippocampi were dissected in cold slice culture dissection medium containing (in mM): 248 sucrose, 26 NaHCO_3_, 10 glucose, 4 KCl, 5 MgCl_2_, 1 CaCl_2_, 2 kynurenic acid and 0.001% phenol red. pH was 7.4, osmolarity 310-320 mOsm kg^-1^, and solution was saturated with 95% O_2_, 5% CO_2_. Tissue was cut into 400 µM thick sections on a tissue chopper and cultured at the medium/air interface on membranes (Millipore PICMORG50) at 37° C in 5% CO_2_. No antibiotics were added to the slice culture medium which was partially exchanged (60-70%) twice per week and contained (for 500 ml): 394 ml Minimal Essential Medium (Sigma M7278), 100 ml heat inactivated donor horse serum (H1138 Sigma), 1 mM L-glutamine (Gibco 25030-024), 0.01 mg ml^-1^ insulin (Sigma I6634), 1.45 ml 5M NaCl (S5150 Sigma)), 2 mM MgSO_4_ (Fluka 63126), 1.44 mM CaCl_2_ (Fluka 21114), 0.00125% ascorbic acid (Fluka 11140), 13 mM D-glucose (Fluka 49152). Wistar rats were housed and bred at the University Medical Center Hamburg-Eppendorf. All procedures were performed in compliance with German law and according to the guidelines of Directive 2010/63/EU. Protocols were approved by the Behörde für Gesundheit und Verbraucherschutz of the City of Hamburg.

#### Anesthetized and awake mice

Male, adult (4-6 months old) C57BL/6J mice were housed and bred in pathogen-free conditions at the University Medical Center Hamburg-Eppendorf. The light/dark cycle was 12/12 h and the humidity and temperature were kept constant (40% relative humidity; 22° C). An AAV2/9 encoding mDlx-postSynTagMA (2.0 × 10^12^ vg ml^-1^), syn-postSynTagMA (3.4 × 10^13^ vg ml^-1^) or syn-GCaMP6f (1.45 × 10^13^ vg ml^-1^) was injected unilaterally in the hippocampus. All procedures performed in mice were in compliance with German law and according to the guidelines of Directive 2010/63/EU. Protocols were approved by the Behörde für Gesundheit und Verbraucherschutz of the City of Hamburg.

### Plasmid construction

To create preSynTagMA, synaptophysin-GCaMP3 was, as previously described (Hoppa et al., 2012), digested by HindIII and BamHI to remove GCaMP3, and CaMPARI was fused to synaptophysin using the In-Fusion® HD Cloning method and kit (Takara Bio USA) after amplifying variants of CaMPARI with PCR using custom primers with base pair overhangs homologous to the synaptophysin plasmid (3’ Primer [CGATAAGCTTTTATGAGCTCAGCCGACC], 5’ Primer [CAGATGAAGCTTATGCTGCAGAACGAGCTTG]). Developing postSynTagMA involved the creation of several intermediate constructs. After removal of restriction site XbaI from pCAG_PSD95.FingR-eGFP-CCR5TC, PSD95.FingR-eGFP-CCR5TC was inserted into a pAAV-hsyn1 backbone without the CAG promoter or upstream zinc finger binding site, to produce pAAV-syn-PSD95.FingR-eGFP-CCR5TC (available upon request). The eGFP was then replaced with CaMPARI1 (Fosque et al., 2015), from which we had deleted the nuclear export signal (NES), to produce pAAV-syn-PSD95.FingR-dNES-CaMPARI1-CCR5TC, a fusion construct of the fibronectin intrabody and a CaMPARI variant that is not restricted to the cytosol and can enter the nucleus. To finally generate pAAV-syn-postSynTagMA, CaMPARI1 was then replaced by CaMPARI2 (without NES and epitope tags) and the point mutations F391W and L398V were introduced using QuickChange PCR to increase calcium affinity (Moeyaert et al., 2018). The left-handed zinc finger (aka CCR5TC) fused to the KRAB(A) transcriptional repressor (Margolin et al., 1994) was removed to produce pAAV-syn-PSD95.FingR-dNES-CaMPARI2_F391W_L398V, the unregulated variant. A sequence including the zinc finger binding sequence (5’-GTCATCCTCATC-3’) (Gross et al., 2013; Perez et al., 2008) upstream of the hsyn1 promoter was synthesized (ThermoFisher) and inserted using the MluI and EcoRI restriction sites to generate pAAV-ZFBS-syn-PSD95.FingR-dNES-CaMPARI2_F391W_L398V-CCR5TC. Other plasmids such as the lower affinity pAAV-syn-PSD95.FingR-dNES-CaMPARI2-CCR5TC, pAAV-syn-synaptophysin-CaMPARI2 and the variants with zinc finger binding sequence are available upon request.

mDlx-postSynTagMA-2A-mCerulean was created by first removing the stop codon from postSynTagMA and inserting a 3’-BsiWI restriction site via PCR. 2A-mCerulean was then inserted 3’of postSynTagMA via Acc65 (5’) and HindIII (3’) to create postSynTagMA-2A-mCerulean. This construct was subsequently cloned via NheI (5’) and BsrGI (3’) into the mDlx-GFP-Fishell1 backbone by replacing the GFP with postSynTagMA-2A-mCerulean to generate mDlx-postSynTagMA-2A-mCerulean.

### Cell culture imaging

SypCaMPARI experiments (**Fig. 1**) were performed at 34° C using a custom-built objective heater. Coverslips were mounted in a rapid-switching, laminar-flow perfusion and stimulation chamber on the stage of a custom-built laser microscope. The volume of the chamber was maintained at ∼75 µl and was perfused at a rate of 400 µl min^-1^. Neurons were perfused continuously during imaging with a standard saline solution containing the following in mM: 119 NaCl, 2.5 KCl, 2 CaCl_2_, 2 MgCl_2_, 25 HEPES, 30 D-Glucose, 10 µM CNQX, and 50 µM D-APV. When noted, 3 µM Tetrodotoxin (TTX) and 20 µM Bicuculline were added to the saline solution.

Neurons were imaged through a EC Plan-Neofluar 40x 1.3 NA objective (Zeiss) or an UAPON40XO340-2 40x 1.35 NA objective (Olympus), using an IXON Ultra 897 EMCCD camera (Andor) at a frame rate of 25 Hz (exposure time: 39.72 ms). Green fluorescence was excited at 488 nm (Coherent OBIS laser, ∼ 3 mW) through a ZET488/10x filter and ZT488rdc dichroic (Chroma). Red fluorescence was excited at 561 nm (Coherent OBIS) through a ZET561/10x filter and ZT561rdc dichroic (Chroma). Green and red fluorescence was collected via ET 525/50m and ET600/50m emission filters (Chroma), respectively.

### Cell culture stimulation and photoconversion

Field stimulation-evoked action potentials were generated by passing 1 ms current pulses, yielding fields of ∼12 V cm^-2^ through the recording chamber bracketed by platinum/iridium electrodes. Electrical stimuli were locked to start according to defined frame number intervals using a custom-built board named “Neurosync” powered by an Arduino Duo chip (Arduino) manufactured by an engineering firm (Sensostar)^51^. A collimated 405 nm LED light source (Thorlabs) was set on top of the microscope stage with a custom-built plastic case (Bob Robertson, Dartmouth College). This light source was coupled to a T-Cube LED Driver (Thorlabs) and a Pulse Pal (Open Ephys) was used to trigger light flashes of specific duration and delay. The light source trigger was set relative to a TTL input from Neurosync. Power density of the 405 nm light was measured using a digital handheld optical power and energy meter with an attached photodiode power sensor (Thorlabs). The power densities used were either 10.8 mW cm^-2^ or 54.1 mW cm^-2^.

For incubator photoconversion experiments (**Supplementary Fig. 1**), custom-built circular 51-diode 405 nm LED arrays (Ultrafire) were wired up to a custom dual programmable relay board to flash light for 100 ms every 10 s, inside a cell culture incubator (New Brunswick; Eppendorf) set to ∼37° C and ∼5% CO_2_. Neurons were then mounted on the microscope with perfusion, and green fluorescence was imaged during 50 stimulations @ 50Hz (as above) to measure calcium-dependent dimming. Only neurons responsive to stimulation were then analyzed (over 90% of cells measured).

#### Experimental setup for variably timed photoconversion

Time locking experiments (**Fig. 2**) were performed on a Nikon Ti-E microscope fitted with an Andor W1 Dual Camera (Andor CMOS ZYLA), dual spinning disk, Coherent Lasers (OBIS 405, 488 and 561 nm) and the Andor Mosaic 3 micro-mirror system, controlled by Andor iQ software, and Nikon elements for image acquisition. PulsePal software was used to time lock the stimulus with the mosaic sequence start. A custom 5×5, 250 µm^2^ square grid was drawn to illuminate different areas (squares) at different times relative to the electrical stimulation. Photoconversion was induced using a train of 5 action potentials (50 Hz) paired with a 100 ms violet light pulse (405 nm, 50 mW cm^-2^) at different delays. The protocol was repeated 5 times at 30 s intervals.

### Cell culture image analysis

EMCCD camera images or confocal image stacks were imported into Fiji. A maximum intensity projection was made from the confocal stacks. Ten pixel diameter circular ROIs were placed over boutons identified by eye using a custom written plugin to localize them over the brightest pixel in the green channel (https://imagej.nih.gov/ij/plugins/time-series.html). Boutons were identified as punctate spots that showed a dimming response to AP stimulation (>98% of punctate spots). ROIs were centered on the brightest green pixel in the green channel and average intensity was measured for red and green channels. Average background fluorescence was determined from several larger ROIs placed across the imaging field where there were no transfected axons. The average green and red background fluorescence was subtracted from the respective values before calculating R/G ratios or ΔF/F.

### Electrophysiology in slice cultures

Hippocampal slice cultures were placed in the recording chamber of the two-photon laser scanning microscope and continuously perfused with an artificial cerebrospinal fluid (ACSF) saturated with 95% O_2_ and 5% CO_2_ consisting of (in mM): 119 NaCl, 26.2 NaHCO_3_, 11 D-glucose, 1 NaH_2_PO_4_, 2.5 KCl, 4 CaCl_2_, 4 MgCl_2_. (pH 7.4, 308 mOsm) at room temperature (21-23° C) or with a HEPES-buffered solution (in mM): 135 NaCl, 2.5 KCl, 10 Na-HEPES, 12.5 D-glucose, 1.25 NaH_2_PO_4_, 4 CaCl_2_, 4 MgCl_2_ (pH 7.4). Whole-cell recordings from CA1 pyramidal neurons were made with patch pipettes (3–4 MΩ) filled with (in mM): 135 K-gluconate, 4 MgCl_2_, 4 Na_2_-ATP, 0.4 Na-GTP, 10 Na_2_-phosphocreatine, 3 sodium-L-ascorbate, and 10 HEPES (pH 7.2, 295 mOsm kg^-1^). In **Supplementary Fig. 3f**, patch pipettes contained (in mM): 135 Cs-MeSO_4_, 4 MgCl_2_, 4 Na_2_-ATP, 0.4 Na-GTP, 10 Na_2_-phosphocreatine, 3 sodium-L-ascorbate, 10 HEPES (pH 7.2, 295 mOsm kg^-1^). Series resistance was below 20 MΩ. A Multiclamp 700B amplifier (Molecular Devices) was used under the control of Ephus ^52^ or Wavesurfer software written in Matlab (The MathWorks).

### Single cell electroporation

At DIV 13-17, CA1 neurons in rat organotypic hippocampal slice culture were transfected by single-cell electroporation^53^. Thin-walled pipettes (∼10 MΩ) were filled with intracellular K-gluconate based solution into which SynTagMA plasmid DNA was diluted to 20 ng µl^-1^. In some experiments, a plasmid encoding mCerulean was also included in the pipette at 20 ng µl^-1^. All experiments were conducted 3-6 days after electroporation. To enable blind analysis of neurons with and without SynTagMA, the electroporation mixes were coded by a second lab member and only after all recordings and analysis were completed was the investigator unblinded. Pipettes were positioned against neurons and DNA was ejected using an Axoporator 800A (Molecular Devices) with 50 hyperpolarizing pulses (−12 V, 0.5 ms) at 50 Hz.

### Recording and analysis of miniature EPSCs and cellular parameters

CA1 pyramidal neurons expressing mCerulean alone or mCerulean plus SynTagMA were patched using only the mCerulean fluorescence to identify them. TTX 1 µM, CPPene 1-10 µM, and picrotoxin 50 µM were added to ACSF (see above). For miniature EPSC (mEPSC) measurements, recording electrodes (3-4 MΩ) contained Cs-gluconate intracellular solution (see above). After the slice rested at least 15 min in the bath, cells were patched and held at −70 mV in the whole-cell voltage clamp configuration (no liquid junction potential correction). EPSCs were recorded 5-15 minutes after break-in. The Event Detection feature of Clampfit 10 (Molecular Devices) was used to detect and measure mEPSC amplitudes and inter-event intervals. For cell parameter measurements, recording electrodes contained K-gluconate solution (see above). Membrane resistance (R_m_) and capacitance (C_m_) was calculated by Ephus using −5 mV voltage steps (50 ms) from a holding potential of −70 mV. Resting membrane voltage was measured in current clamp mode. A 1 s current ramp (0 to +600 pA) was injected to measure the rheobase and the voltage threshold for action potentials. Firing rates were calculated from 1 s current steps (−400 pA to +600 pA).

### Two-photon microscopy in hippocampal slices

Two-photon imaging was performed in rat organotypic hippocampal slice cultures and mouse acute hippocampal slice. The custom-built two-photon imaging setup was based on an Olympus BX51WI microscope equipped with LUMPlan W-IR2 60× 0.9 NA (Olympus), W Plan-Apochromat 40× 1.0 NA (Zeiss) or IRAPO 25x 1.0 NA (Leica) objectives controlled by the open-source software package ScanImage^54^. Two pulsed Ti:Sapphire lasers (MaiTai DeepSee, Spectra Physics) controlled by electro-optic modulators (350-80, Conoptics) were used to excite SynTagMA green (980 nm) and red species (1040 nm), respectively. When Z-stacks of SynTagMA-expressing neurons were acquired, each plane was scanned twice, using 980 nm and 1040 nm excitation, respectively. For quantification of pre- or postsynaptic targeting, axons of CA3 cells or oblique dendrites of CA1 neurons expressing mCerulean and SynTagMA were imaged at 840 nm and 980 nm. For three-color experiments, separate stacks were taken to image mCerulean (840 nm) and SynTagMA (980 nm & 1040 nm). Emitted photons were collected through the objective and oil-immersion condenser (1.4 NA, Olympus) with two pairs of photomultiplier tubes (H7422P-40, Hamamatsu). 560 DXCR dichroic mirrors and 525/50 and 607/70 emission filters (Chroma) were used to separate green and red fluorescence. Excitation light was blocked by short-pass filters (ET700SP-2P, Chroma). Two brief violet light pulses (395 nm, 100 ms, 16 mW mm^-2^, 0.1 Hz) were delivered through the objective using a Spectra X Light Engine (Lumencor) just before imaging to photoswitch the CaMPARI moiety into its bright state^10^.

### Spine density measurement and analysis

Two-photon microscopy at 840 nm was used to excite cells expressing mCerulean or mCerulean plus SyntagMA as described above. An Olympus LUMFL N 60x 1.1NA objective (PSF: 0.35 x 0.35 x 1.5 µm) was used to collect Z-stacks (0.3 µm z-step) of *stratum radiatum* proximal oblique dendrites (∼100 µm from soma). Image stacks were deconvolved using a blind deconvolution algorithm in Autoquant X3 (Media cybernetics). Spines and dendrites were semi-automatically detected using the Filament tracer feature of Imaris (Oxford Instruments). For each neuron, spine density was determined from 1-3 dendrites of 30-100 µm length (**Supplementary Fig. 3g**).

### Photoconversion in hippocampal slices

Photoconversion was achieved by delivery of violet light (395 nm, 16 mW mm^-2^, duration 100 ms – 2 s as indicated in figure legends) using a Spectra X Light Engine (Lumencor) coupled with a liquid light guide to the epifluorescence port of the two-photon microscope. During the violet light pulses, shutters (Uniblitz) protected the photomultiplier tubes. We typically used 100-500 ms violet light pulses repeated 15-50 times with a 1 s delay from stimulus onset to photoconvert postSynTagMA. Image stacks were acquired prior to and following photoconversion.

### Photoconversion of back-propagating action potentials

The first image stack was acquired before patching the SynTagMA-expressing neuron. The cell was then whole-cell patch-clamped and bAPs (100 Hz) were evoked by somatic current injection and paired with 500 ms photoconversion light (1 s delay) in the presence of CPPene (10 µM) and NBQX (10 µM) to block synaptic transmission. This pairing was repeated 15 times. While acquiring the second image stack, the cell was held in voltage clamp at −65 mV to prevent depolarization. The field of violet light illumination for these experiments was 557 um in diameter and therefore large enough to illuminate the entire dendritic arbor.

### Propidium iodide staining

Organotypic hippocampal slices aged between DIV 18-28 were used. 2-4 days before staining, CA1 neurons were electroporated with mCerulean (20 ng µl^-1^) and postSynTagMA (20 ng µl^-1^). Slices were placed in the two-photon imaging chamber in HEPES-buffered solution containing 3 µM propidium iodide (PI). The objective was then placed above the CA1 region of the slice and illuminated with violet light (395 nm, 16mW mm^-2^). Following illumination, slices were kept in the PI solution for 30-45 minutes and subsequently imaged (840 nm excitation). PI fluorescence was collected through 607/70 filters and mCerulean fluorescence was collected through 525/50 filters. NMDA excitotoxicity was used as a positive control. Slices were exposed to 1mM NMDA in HEPES-buffered solution for 1.5 hours and then labeled with PI for 30-45 minutes before imaging as above. To quantify violet light toxicity, PI-positive nuclei were counted in each 2P image stack (background subtracted, median filtered) and normalized to the tissue volume.

### Quantification of pre- and postSynTagMA localization

A macro written in Fiji^21^ was used for two-photon 3D image analysis at 840, 980 and/or 1040 nm wavelengths. When z-stacks contained alternating images collected at different excitation wavelengths, they were first separated and then xyz-aligned to correct for chromatic aberration using green and/or red channels and the pairwise stitching plugin^55^. mCerulean and SynTagMA fluorescence values were obtained from images after median filtering and rolling ball background subtraction^56^. Regions of interest (ROI) were drawn onto maximum intensity projections and compared to axonal or dendrite shafts, respectively. Only spines projecting laterally from the dendrite were analyzed.

### Semi-automatic segmentation and analysis of SynTagMA

For the initial characterization of postSynTagMA, quantification of green to red conversion was performed manually in a small sets of synapses (<50) in Matlab and/or Fiji. Covering relative larger areas of the dendritic tree gave rise to larger data sets (>1000 synapses) that made a semi-automated analysis approach necessary. Rigid registration was performed using the Pairwise stitching plugin^55^ in Fiji. The broad emission spectrum of autofluorescent objects tuned out to be very useful to align the red channel from the 980 nm stack to the green channel from the 1040 nm stack. Image stacks were further processed using a blind deconvolution algorithm in Autoquant X3. Particularly critical was to correct for position change of synapses in 3D due to tissue deformation (“warping”) between acquisitions of image stacks (pre- and post-stimulation). To correct for tissue warping, synapse voxels were reassigned to their initial position using either elastix^23^ or ANTs (Advanced Normalization Tools^57^). To detect synapses, we used Imaris 3D spot detection on the deconvolved G_0_ signal. Spots (volumetric ROIs) were used to extract fluorescence intensities from green and red channels (maximum pixel values from non-deconvolved, median-filtered images) at every time point. To quantify photoconversion, green (G_0_ and G_1_) and red (R_0_ and R_1_) values were normalized to the population mean of G_0_ and R_0_, respectively.

### Automated segmentation and analysis of SynTagMA

To analyze thousands of synapses in large 3D datasets, we developed a pipeline to automatically identify spots in 3D and define volumes (ROIs) for accurate measurements of fluorescence changes. After two microscope-specific pre-processing steps, the data is processed by SynapseLocator, a GUI-controlled software package written in Matlab, which calls specific subroutines (Fiji, elastix, Weka) to perform the analysis:

1. De-interleaving of 980/1040 nm image stacks to separate green and red channels
2. Rigid registration to correct for chromatic aberration and beam misalignment
3. Deconvolution and registration of post-image to pre-image using elastix
4. Random forest model, built with machine learning
5. Spot identification, size checks, ROI creation
6. Calculation of SynTagMA photoconversion in ROIs

A specific challenge was posed by local movements of individual synapses between the time points of data acquisition, which made *post hoc* alignment and de-warping of images an essential step prior to segmentation. A typical experiment consisted of at least two time points (before / after stimulation) in which green and red fluorescence was collected near-simultaneously, alternating between 980 and 1040 nm excitation at every optical section of the z-stack. For each time point, we corrected channel misalignment due to chromatic aberration by performing rigid registration of the de-interleaved two-color images in Matlab. The detailed workflow in SynapseLocator consists of an initial deconvolution step (Fiji, DeconvolutionLab2 plugin^22^) in which diffraction-induced blurring of the images was reduced. We used elastix^23^ to register image stacks across time points (only green channel; G_0_ and G_1_). Registration proceeds in four steps (rigid transformation, rotation transformation, affine transformation, and non-rigid registration) for which we provide optimized parameter sets. The transformation was then applied to both red and green channel data (deconvolved and raw). Synapse detection involved a machine learning process accessed via Fiji (Weka, Trainable Weka Segmentation plugin^24^). Two classes are manually identified (“spot”, “no spot”) to train a random forest model considering a set of scale-invariant image features (Hessian and Laplacian, each calculated at three scales, Fiji, FeatureJ plugin). Typically, the model was robust enough to allow for analysis of all experiments from a series. A synapse is identified as a region in which a group of voxels with spot class properties shows minimal connectivity (26-connected neighborhood). For each region, an ellipsoid enclosing the identified connected voxels is calculated in Matlab and stored as ROI. Regions are further filtered by size to exclude objects with extreme values. Inclusion criteria for synapses are: green initial signal (G_0_) above background, red initial signal (R_0_) below user-defined threshold, high shape correlation between G_0_ and R_1_. Converted synapses are identified according to their “conversion-factor” (ΔR / (G_0_+G_1_)).

### Distance-dependence of bAP-induced calcium transients

CA1 neurons expressing mCerulean and postSynTagMA were used for these experiments. Prior to patching, z-stacks were acquired along the apical dendritic tree to image postSynTagMA (980 nm / 1040 nm). Two to three stacks were required to image the apical dendritic tree. The time taken to image all stacks was 30-40 minutes. The cell was then whole-cell patch-clamped and 50 bAPs were paired with 500 ms violet light (1 s delay). This pairing was repeated 15 times (0.1 Hz). Immediately after, all 980/1040 nm image stacks were acquired followed by complementary stacks of mCerulean at 840nm for morphology.

In CA1 neurons expressing jGCaMP7b, frame scans (6 x 6 µm) of oblique dendrites along the entire dendritic arbor were acquired (980 nm) while evoking 50 bAPs. In each trial, 50 frames (64 x 64 pixels) were acquired at 5.9 Hz. At least 3 trials were recorded from each section of dendrite. Following calcium imaging, the morphology of the entire cell was imaged using jGCaMP7b baseline fluorescence. The fluorescence time course was measured by placing ROIs on individual spines in Fiji. We calculated ΔF/F_0_; F_0_ was determined 300 ms prior to bAP onset. To analyze photoconversion (postSynTagMA) or calcium transients (jGCaMP7b) in synapses as a function of distance from the soma, we created a series of 20 µm wide concentric rings around the soma using a custom-written MATLAB script. For the synapses located in each ring, we calculated the median photoconversion (or median calcium transient amplitude) and interquartile range.

### postSynTagMA turnover measurements

postSynTagMA turnover experiments were performed on dissociated hippocampal neurons (DIV 14-16, 9-11 days post-transfection). Cells were imaged at ∼21°C in a modified Tyrode’s solution containing the following (in mM): 119 NaCl, 2.5 KCl, 3 CaCl2, 1 MgCl2, 25 HEPES, and 30 D-Glucose with 10 µM CNQX. PostSynTagMA photoconversion was induced using a train of 50 action potentials (50 Hz) paired with a 500 ms photoconverting light pulse in custom-drawn rectangular regions (18.6 mW mm^-2^, 500-750 ms delay). Z-stacks with 0.25 µm step size were collected over the course of an hour every 5-6 minutes. Cells were illuminated at each step by 405nm (1% intensity), 488 (25% intensity) and 561 nm (20% intensity) lasers.

### Photoconverting sub- and suprathreshold responses

For extracellular synaptic stimulation, a monopolar electrode was placed in *stratum radiatum* and two 0.2 ms pulses, 40 ms apart, were delivered using an ISO-Flex stimulator (A.M.P.I.). Stimulation intensity was adjusted to be subthreshold for action potentials (i.e. to evoke ∼15 mV EPSPs or ∼-500 pA EPSCs) or suprathreshold (i.e. evoking action potentials) by patching a nearby neuron in CA1. Stimulation was paired with 100 ms violet light (1 s delay) 50 times.

### Subthreshold optogenetic stimulation and photoconversion

An AAV2/9 synapsin-ChR2(ET/TC)-2A-synaptophysin-mCerulean, prepared at the UKE vector facility, was injected locally into CA3 of slice cultures at DIV 7. Additionally, several CA1 neurons were electroporated with postSynTagMA (20 ng µl^-1^) at DIV 22. CA3 neurons were stimulated with blue light pulses (470 nm, 2 pulses, 2-5 ms duration, 40 ms apart) applied through a light fiber placed above the CA3. The light intensity and pulse width were set to evoke EPSCs (0.5 – 1.0 nA) in a neighboring CA1 neuron. Stimulation was paired with 100 ms violet light (1 s delay) and repeated 50 times at 0.1 Hz to photoconvert postSynTagMA in active spines. Z-stacks of spiny dendrites were acquired before and after photoconversion. To image mCerulean-labeled boutons, a third z-stack at 840nm was acquired immediately after the post-photoconversion stack.

### Relabeling postSynTagMA

For relabeling experiments, paired-pulse extracellular stimulation (0.2 ms pulses, 40 ms apart) was combined with 250 ms of violet light. This was repeated 25 times (0.1Hz) and a second image stack was acquired immediately thereafter. After imaging, the slice was returned to the incubator. Approximately 18 h later, the procedure was repeated.

### Virus injection and hippocampal window surgery

Mice were anesthetized with an intraperitoneal injection of Ketamine/Xylazine (0.13/0.01 mg g^-1^ bodyweight) and placed on a heating blanket to maintain the body temperature. In addition, mice received a subcutaneous dose of Carprofen (4 mg kg^-1^) for post-surgery analgesia. Eyes were covered with eye ointment (Vidisic, Bausch + Lomb) to prevent drying. Prior to surgery, the depth of anesthesia and analgesia was evaluated with a toe-pinch to test the paw-withdrawal reflex. Subsequently, mice were fixed in a stereotactic frame, the fur was removed with a fine trimmer and the skin of the head was disinfected with Betaisodona solution using sterile cotton swabs. The skin was removed by a midline scalp incision (1-3 cm), the skull was cleaned using a bone scraper (Fine Science Tools) and a small hole was drilled with a dental drill (Foredom) above the injection site. AAV2/9-mDlx-SynTagMA-2A-mCerulean or AAV2/9-syn-GCaMP6f was targeted unilaterally to the dorsal CA1 area (−2.0 mm AP, ± 1.3 mm ML, - 1.5 mm DV relative to Bregma. 0.6 µl of virus suspension was injected. All injections were done at 100 nl min^-1^ using a glass micropipette. After the injection, the pipette stayed in place for at least 5 min after virus delivery before it was withdrawn and the scalp was closed with sutures. During the two days following surgery animals were provided with Meloxicam mixed into soft food.

Two weeks after virus injection, mice were anesthetized as described above to implant the hippocampal window. After fur removal, skin above the frontal and parietal bones of the skull was removed by one horizontal cut along basis of skull and two rostral cuts. The skull was cleaned after removal of the periosteum, roughened with a bone scraper and covered with a thin layer of cyanoacrylate glue (Cyano Veneer). After polymerization a 3-mm circle was marked on the right parietal bone (anteroposterior, −2.2 mm; mediolateral, +1.8 mm relative to bregma) with a biopsy punch and the bone was removed with a dental drill (Foredom). The bone fragment and dura were carefully removed with fine surgical forceps. The cortex above the hippocampus was aspirated with a 0.8 mm blunt needle connected to a water jet pump. When first layer of fibers running orthogonal to midline became visible, a 0.4 mm blunt needle was used to carefully remove the upper two of three fiber layers of the external capsule. Absorbable gelatin sponges (Gelfoam, Pfizer) were used to control bleeding. The craniotomy was washed with sterile PBS and a custom-built imaging window was inserted over the dorsal hippocampus. The window consisted of a hollow glass cylinder (diameter: 3 mm, wall thickness: 0.1 mm, height: 1.5 mm) glued to a No. 1 coverslip (diameter: 3mm, thickness: 0.17 mm) on the bottom and to a stainless steel rim on the top with UV-curable glass glue (Norland NOA61). The steel rim and a head holder plate (Luigs & Neumann) were fixed to the skull with cyanoacrylate gel (Pattex). After polymerization, cranial window and head holder plate were covered with dental cement (Super Bond C&B, Sun Medical) to provide strong bonding to the skull bone. During the two days following surgery animals were provided with Meloxicam mixed into soft food.

### Hippocampal imaging and photoconversion *in vivo*

Mice were anesthetized with isoflurane at a concentration ranging between 2.0% and 2.5% in 100% O_2_, head-fixed under the microscope on a heated blanket to maintain body temperature. Eyes were covered with eye ointment (Vidisic, Bausch + Lomb) to prevent drying. The window was centered under the two-photon microscope (MOM-scope, Sutter Instruments, modified by Rapp Optoelectronics) using a low-magnification objective (4x Nikon Plan Fluorite) and reporter expression (postSynTagMA or GCaMP6f) was verified in the hippocampus using epi-fluorescence. The cranial window was then filled with water to image in two-photon mode through a 40x objective (40X Nikon CFI APO NIR, 0.80 NA, 3.5 mm WD). The green species of SynTagMA or GCaMP6f was excited with a Ti:Sa laser (Chameleon Vision-S, Coherent) tuned to 980 nm. The red species of SynTagMA was excited with an ytterbium-doped 1070-nm fiber laser (Fidelity-2, Coherent). Single planes (512×512 pixels) were acquired at 30 Hz with a resonant-galvanometric scanner at 29-60 mW (980 nm) and 41-60 mW (1070 nm) using ScanImage 2017b (Vidrio). Emitted photons were detected by a pair of photomultiplier tubes (H7422P-40, Hamamatsu). A 560 DXCR dichroic mirror and 525/50 and 607/70 emission filters (Chroma Technology) were used to separate green and red fluorescence. Excitation light was blocked by short-pass filters (ET700SP-2P, Chroma). Since SynTagMA fluorescence was relatively low, 450 - 1800 frames per optical plane were acquired and averaged. For identification of active CA1 neurons (**Fig 7A-C**), the conversion protocol consisted of 2 s violet light pulses (405 nm, 12.1 mW mm^-2^) repeated 10 times in awake and ketamine/xylazine anesthetized mice. In awake behaving mice, photoconversion light was triggered upon water reward delivery. Under isoflurane anesthesia (**Fig. 7G**), conversion of active synapses in interneurons was achieved with 20 flashes of 3 s violet light pulses (405 nm, 0.42 mW mm^-2^). Motion artefacts were corrected with a custom-modified Lucas-Kanade-based alignment algorithm in Matlab.

## QUANTIFICATION AND STATISTICAL ANALYSIS

Statistical analysis was performed with GraphPad Prism (v8) or Matlab. For data with normal distributions, Student’s t-test or one-way ANOVA were used. Data considered non-normal (according to a D’Agostino-Pearson test) underwent non-parametric tests (Kruskal-Wallis).

