## Supplementary Figures for "Freeze-frame imaging of synaptic activity using SynTagMA"

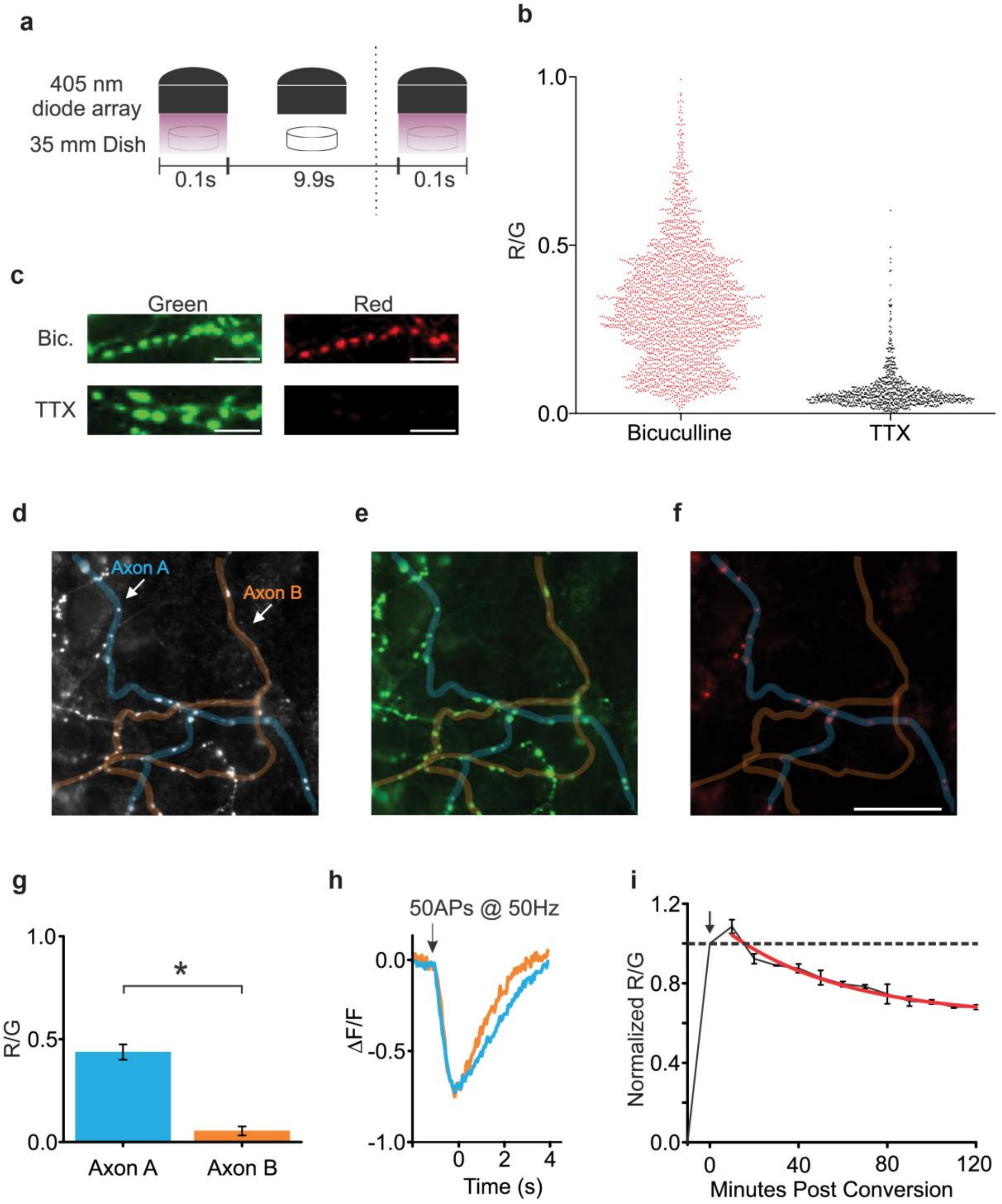


**Supplementary Figure 1: Spontaneous circuit activity of cultured primary hippocampal neurons drives photoconversion of preSynTagMA**.

**(a)** LED arrays (51 diodes, 405 nm) were placed in an incubator (5% $CO_2$, 37°C) and set to flash for 100 ms every 10 s for 4 hours. LED arrays were placed on top of 35 mm dishes containing cultured primary hippocampal neurons. **(b)** R/G ratio of individual boutons incubated with bicuculline (3 dishes, 2,456 synapses) or TTX (4 dishes, 866 synapses) while illuminating with violet light as in a. **(c)** Example green and red raw fluorescence of boutons from b). **(d)** Axons from two neurons in bicuculline condition (axon A, blue overlay; axon B, orange overlay). **(e)** Raw green fluorescence. **(f)** Raw red fluorescence. Note that photoconversion (red fluorescence) was restricted to boutons of axon A. **(g)** Mean R/G ± SE from boutons of axon A (n = 11 boutons, R/G = 0.44 ± 0.04) and axon B (n = 12 boutons, R/G = 0.05 ± 0.02) (Student's t-test, unpaired, *p < 0.001), indicating differential activity. **(h)** Dimming of green fluorescence from axons

A (blue) and B (orange) during a 50 AP, 50 Hz stimulus train. Similar dimming indicates both axons could fire action potentials and had similar calcium influx during induced spiking. Therefore, the lack of photoconversion in bicuculline was most likely due to a lack of spiking in axon B. **(i)** Slow decay of R/G ratio following strong photoconversion (40 x 1 s 405 nm light, paired exactly with 20 APs @ 50Hz). Images were acquired every ten minutes for two hours (n = 4 cells). Data are shown as mean ± SEM.

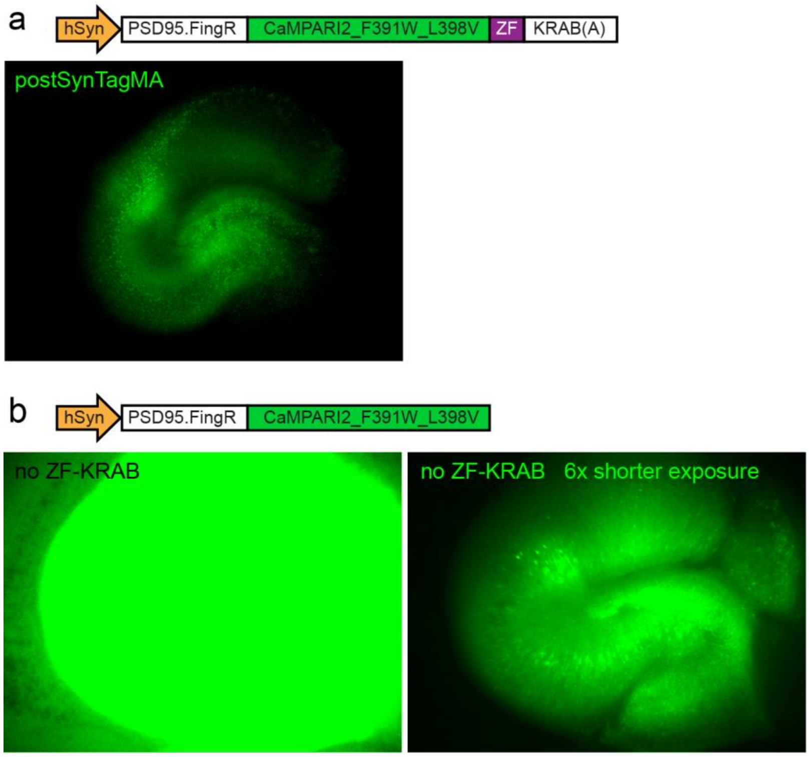


**Supplementary Figure 2: Effect of autoregulatory elements on postSynTagMA expression level.**

**a)** Organotypic rat hippocampal slice culture 4 days after AAV9-mediated transduction with postSynTagMA. **(b)** Left: Same as a), but transduced with PSD.95-FingR_CaMPARI2_F391W_L398V (no ZF-KRAB). Virus titers and applied volume were matched, as were the illumination intensity and camera settings. Right: Same culture, image was de-saturated by six-fold reduction in exposure time.

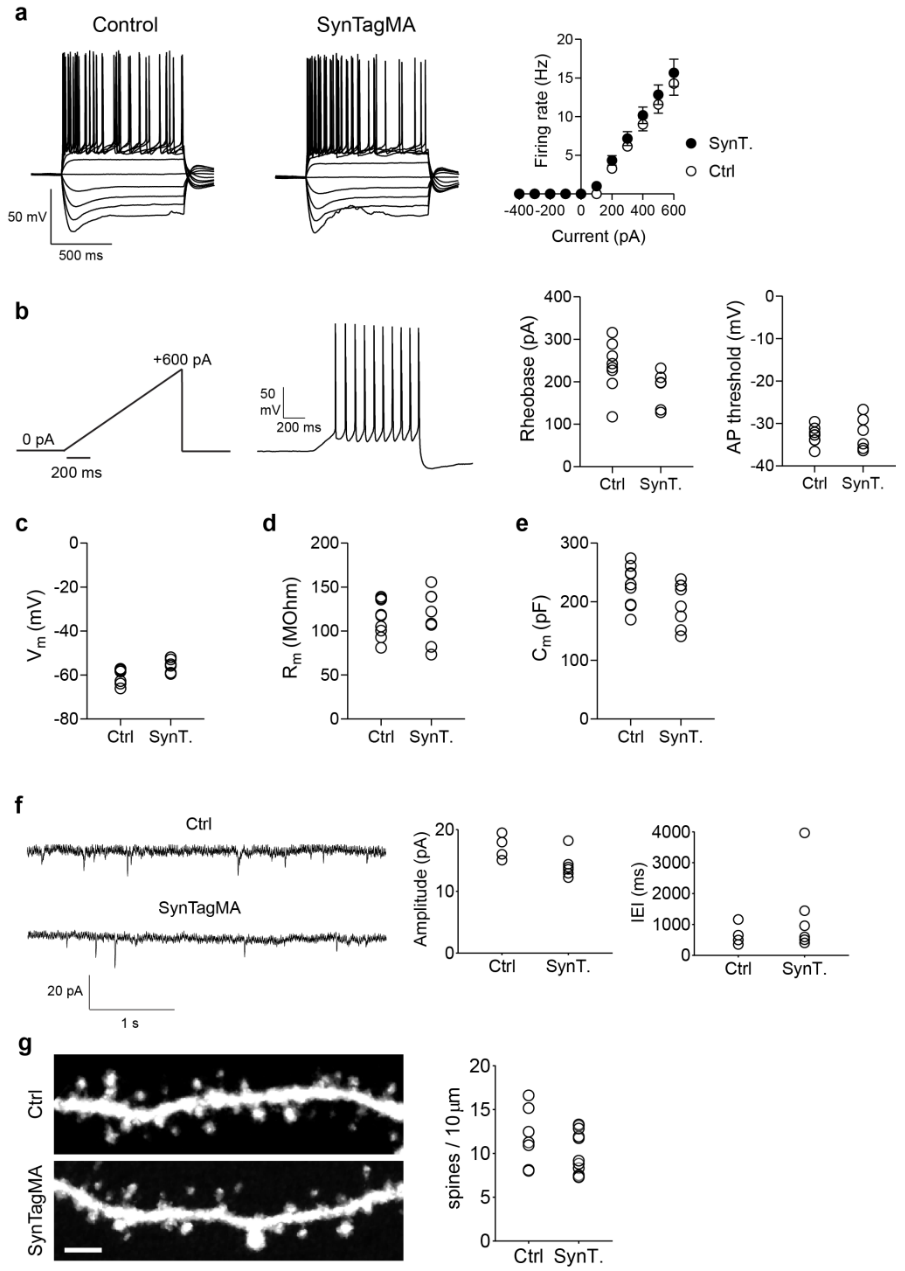


**Supplementary Figure 3: Active and passive properties, synaptic inputs and spine density of SynTagMA-expressing neurons**.

**(a)** Voltage responses of CA1 pyramidal neurons expressing either mCerulean (Ctrl) or mCerulean and postSynTagMA (SynTagMA) elicited by increasing current steps from -400 pA to +600 pA (1 s duration at 0.067 Hz). Action potential firing rates increased linearly for depolarizing current injections. **(b)** Current ramp (1 s duration from 0 to +600 pA) and example response of a SynTagMA-expressing neuron used to determine the rheobase and action potential (AP) threshold. Rheobase of SynTagMA-expressing neurons was not different from mCerulean controls (Ctrl: 235.2 ± 21.4 pA, n = 8; SynT: 183.2 ± 17.4 pA, n = 6; P = 0.108, Mann Whitney). AP threshold of SynTagMA-expressing neurons was not different from mCerulean controls (Ctrl: -32.8 ± 0.74 mV, n = 8; SynT: -32.4 .± 1.6 mV, n = 6, P = 0.80, Student's t test). **(c)** Resting membrane potential (Vm, Ctrl: -59.9 ± 1.1 mV, n = 9; SynT: -55.7 ± 1.3 mV, n = 6; P = 0.088, Mann Whitney). **(d)** Membrane resistance (Rm) (Ctrl: 114.2 ± 6.9 MΩ, n = 9; SynT: 115.8 ± 10.2 MΩ; n = 8, P = 0.90, Student's t test). **(e)** Membrane capacitance (Cm) (Ctrl: 227 ± 12 pF, n = 9; SynT: 197 ± 13 pF, n = 8; P = 0.105, Student's t test). **(f)** Representative miniature AMPA receptor-mediated postsynaptic currents (mEPSCs) from CA1 pyramidal cells voltage clamped at -70 mV in the presence of CPPene, picrotoxin and tetrodotoxin. Mean mEPSC amplitude was 17.1 ± 1.0 pA for control neurons (n = 4) and 14.2 ± 0.9 pA (n = 6) for SynTagMA-expressing neurons (P = 0.067, Mann Whitney). Inter-event intervals were 667 ± 176 ms for control neurons (n = 4) and 1314 ± 553 ms (n = 6) for SynTagMA-expressing neurons (P = 0.48, Mann Whitney test). **(g)** Maximum intensity projection of deconvolved two photon images from CA1 neurons expressing mCerulean alone (above) or mCerulean and SynTagMA (below). Scale bar is 5 µm. Mean spine density per 10 microns (± SEM) was 11.8 ± 1.2 for mCerulean-expressing neurons (7 cells, 3 slices, 2896 spines) and 10.1 ± 0.7 for mCerulean and SynTagMA-expressing neurons (11 cells, 5 slices, 3946 spines; P = 0.42, Mann Whitney).

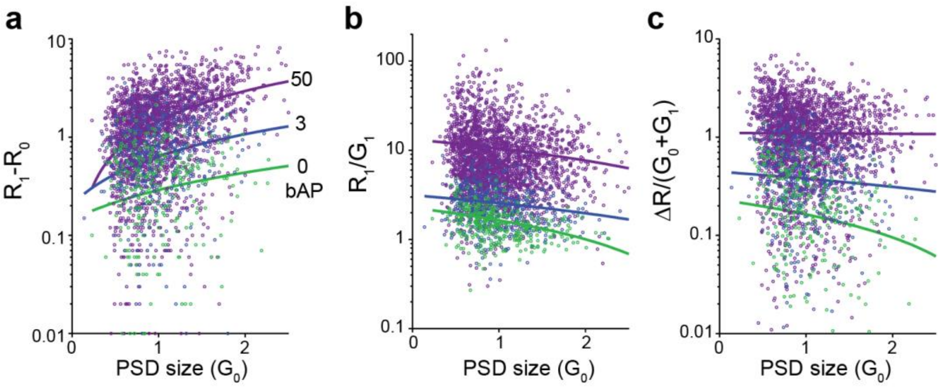


**Supplementary Figure 4: Comparison of conversion metrics vs synapse size (a)** Absolute change in red fluorescence $\Delta R = R_1 - R_0$, plotted against PSD size ($G_0$). Lines are linear regression on semi-log plot. (0 bAPs: $r^2$ = 0.018, slope = 0.146; 3 bAPs: $r^2$ = 0.049, slope = 0.435; 50 bAPs: $r^2$ = 0.170, slope = 1.50). **(b)** Using only the time point after photoconversion. R1/G1 versus PSD size (0 bAPs: $r^2$ = 0.12, slope = -0.63; 3 bAPs: $r^2$ = 0.026, slope = -0.59; 50 bAPs: $r^2$ = 0.010, slope = -2.8) **(c)** $(R_1 - R_0)/(G_1 + G_0)$ versus PSD size (0 bAPs: $r^2$ = 0.010, slope = -0.068; 3 bAPs: $r^2$ = 0.0029, slope = -0.066; 50 bAPs: $r^2$ = $1.77 \times 10^{-5}$, slope = -0.011). The X axis is truncated at 2.5 and Y axis at the lower end (0.01).

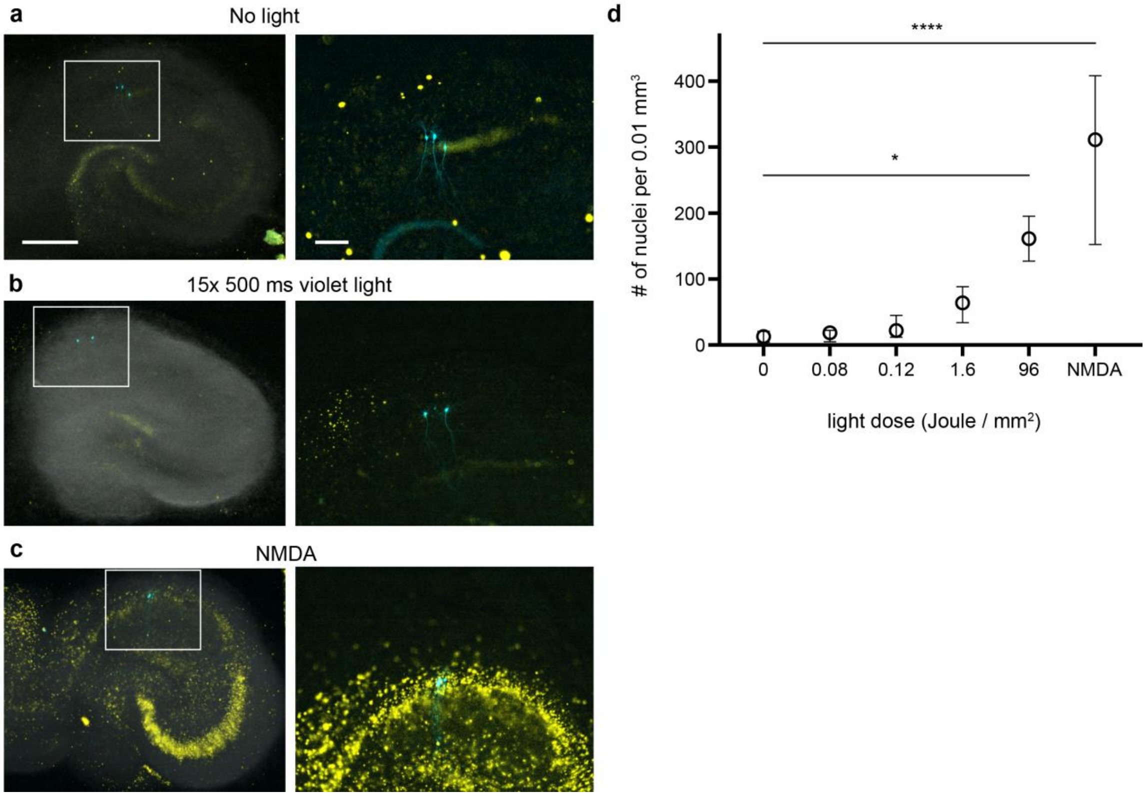


**Supplementary Figure 5: Assessing photodamage caused by violet light and NMDA in organotypic cultures by propidium iodide (PI) staining**.

**(a-c)** Representative images of hippocampal slice cultures following incubation with propidium iodide, which enters and stains damaged cells. Overlay of Dodt-contrast image (grayscale), propidium iodide fluorescence (yellow) and cerulean-expressing CA1 pyramidal cells (cyan). Scale bars 500 µm and 200 µm, respectively. **(a)** Slice culture kept in the dark. **(b)** After exposure to our typical photoconversion protocol (395 nm 16 mW $mm^{-2}$, 15 x 500 ms, 0.12 J $mm^{-2}$). **(c)** After exposure to 1 mM NMDA for 1 hour. **(d)** Quantification of results vs light dose. Plotted are median and interquartile range. * $p < 0.0461$, **** $p < 0.0001$, a one-way ANOVA followed by Dunnett's multiple comparisons to the no light condition. No light (n = 5 slices), 0.08 Joule (n = 3 slices), 0.12 Joule (n = 5 slices), 1.6 Joule (n = 3 slices), 96 Joule (n = 2 slices), NMDA (n = 4 slices).

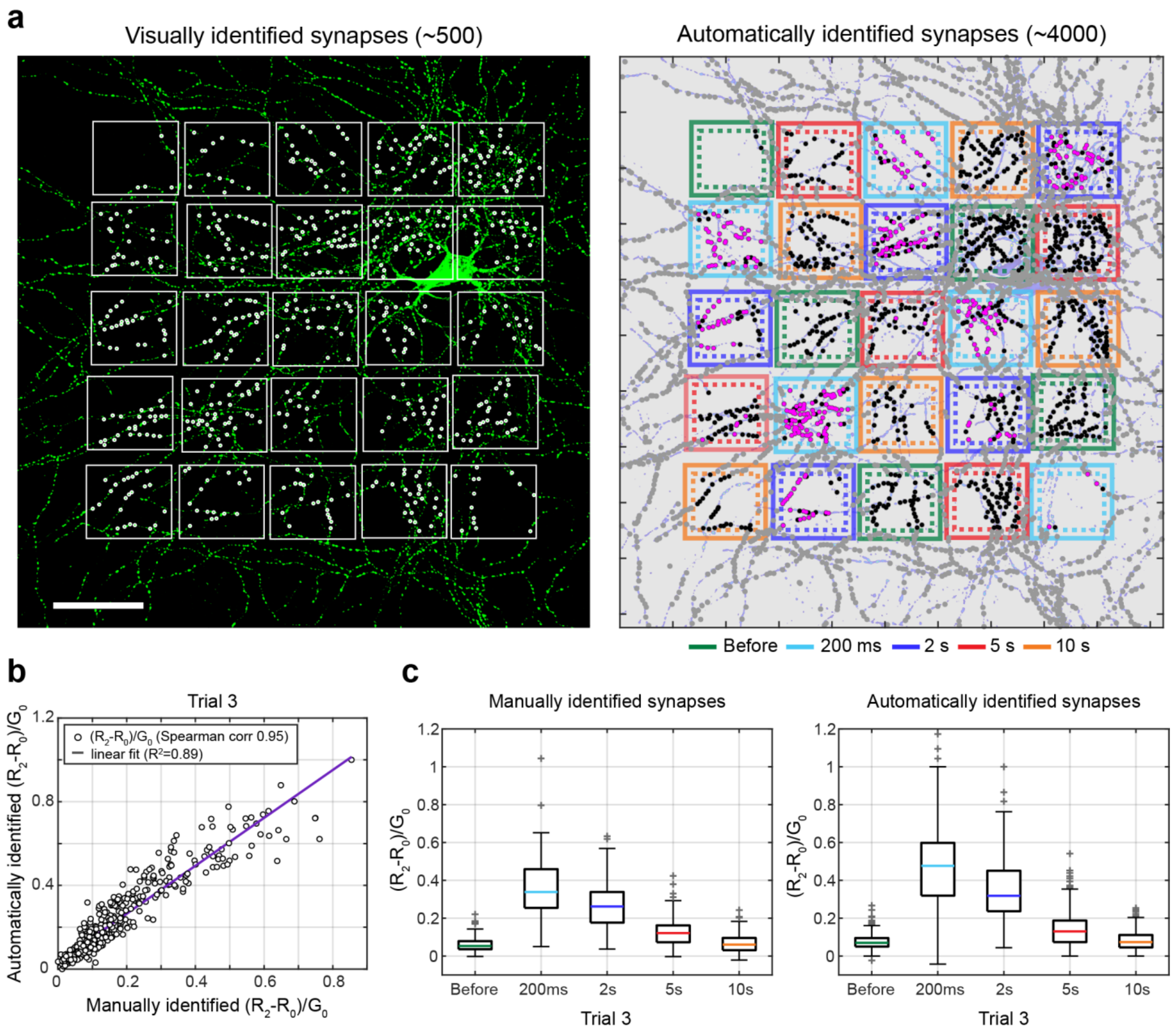


**Supplementary Figure 6: Comparison of analysis results with manual identification of ROIs vs analysis performed with SynapseLocator**.

**(a)** Full-field view (maximum projection) of the same cultured hippocampal neuron expressing preSynTagMa as shown in Fig. 2 with overlayed photoconversion grid. Dashed lines are borders set for including automatically identified boutons in a group. Left: Regions of interest (~500 ROIs) were selected and manually curated using Fiji (white dots are ROIs). Right: Colored and grey dots indicate (ROIs) identified and analyzed by SynapseLocator. Grey spots were outside the illuminated fields and not quantified. Magenta spots: $(R_2 - R_0)/G_0 > 0.25$. Black spots: $(R_2 - R_0)/G_0 < 0.25$. Scale bar is 50 µm. **(b)** The photoconversion values from Trial 3 (see Fig. 2) from the group of boutons that were both manually and automatically identified plotted against each other (n = 413 boutons). **(c)** Photoconversion $(R_2 - R_0)/G_0$ vs timing delay for the same data manually identified (right, n = 583 boutons) or identified and analyzed using SynapseLocator (left, n = 1059 boutons). Boxes: Median, 25% and 75% percentiles and whiskers the 1.5 interquartile interval. Grey + are outliers. The data distribution was not normal within any group (D'Agostino & Pearson test, P < 0.0001). A non-parametric Kruskal-Wallis test followed by Dunn's multiple comparisons (before manual vs before automatic, exact manual vs exact automatic etc.) showed the outcome was not significantly different at any time point.

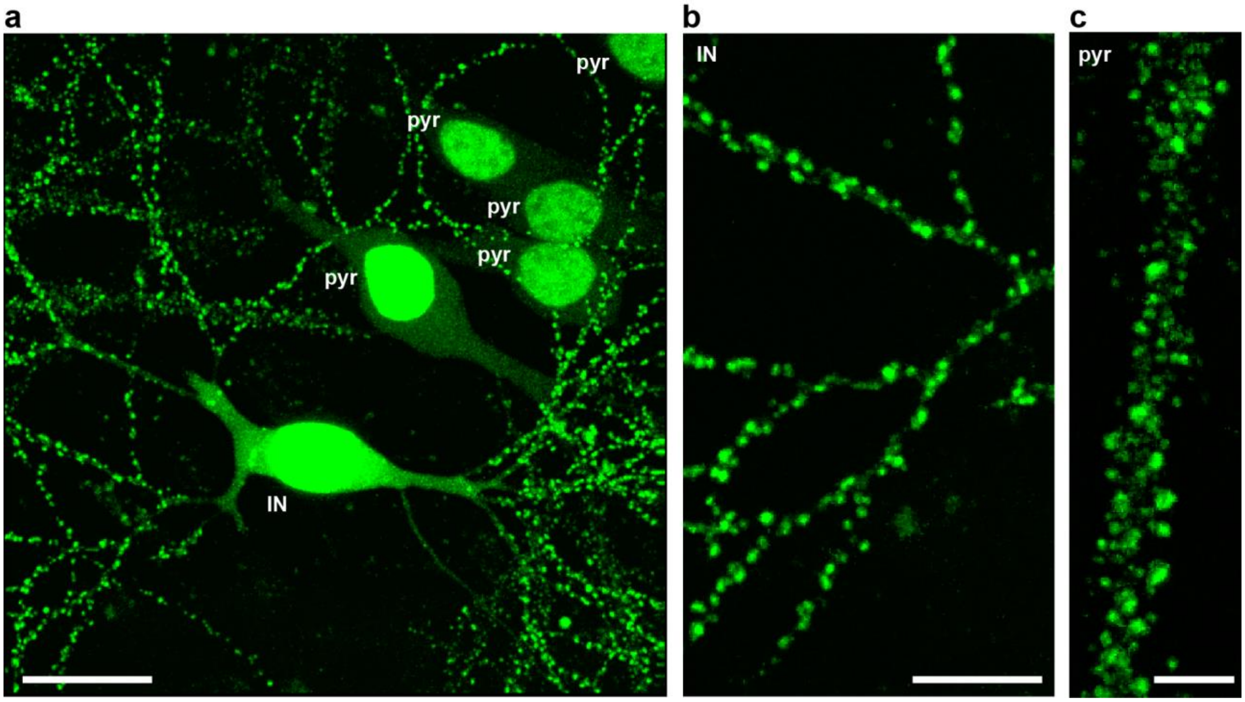


**Supplementary Figure 7. PostSynTagMA identifies excitatory synapses on interneurons**.

**(a)** Two-photon image (maximum intensity projection) showing expression of postSynTagMA in an interneuron (IN) next to five pyramidal CA1 cells (pyr). Scale bar 20 µm. **(b)** Detail of postSynTagMA puncta in smooth interneuron dendrite (IN) and **(c)** spiny pyramidal neuron dendrite (pyr). Scale bars a) 20 µm b) 10 µm c) 5 µm.

### SUPPLEMENTARY MOVIES

#### Supplementary Movie 1: Spine in contact with Channelrhodopsin-expressing bouton gets photoconverted

3D video showing automated detection of synapses (silver objects) based on green fluorescence of postSynTagMA, red fluorescence inside the detected objects (masked), and zoom-in to a strongly photoconverted synapse (true positive) in contact with a presynaptic bouton (cyan). Note the large number of non-photoconverted synapses that are also distant from activated presynaptic terminals (true negatives).

#### Supplementary Movie 2: In vivo calcium imaging of CA1 neurons expressing GCaMP6f

Calcium imaging reveals similar patterns of activity during awake and under isoflurane anesthesia, but not under ketamine/xylazine anesthesia. Different conditions were tested on the same animal and CA1 area on different days. Ketamine/xylazine anesthesia reduced the number of active neurons and the intensity of calcium transients.

#### Supplementary Movie 3: In vivo calcium imaging of CA1 neurons using postSynTagMA

Nuclear calcium elevations during treadmill running are visible as dimming of postSynTagMA green fluorescence. Total time: 4 min.
